# A Critical Role for the Fascin Family of Actin Bundling Proteins in Axon Development, Brain Wiring and Function

**DOI:** 10.1101/2025.02.21.639554

**Authors:** Katherine R. Hardin, Arjolyn B. Penas, Shuristeen Joubert, Changtian Ye, Kenneth R. Myers, James Q. Zheng

**Author notes:** Correspondence: James Zheng, PhD, Department of Cell Biology, 615 Michael Street, Atlanta, Georgia 30322, USA.

## Abstract

Actin-based cell motility drives many neurodevelopmental events including guided axonal growth. Fascin is a major family of F-actin bundling proteins, but its role in axon development *in vivo* and brain wiring remains unclear. Here, we report that fascin is required for axon development, brain wiring and function. We show that fascin is enriched in the motile filopodia of axonal growth cones and its inhibition impairs axonal extension and branching of hippocampal neurons in culture. We next provide evidence that fascin is essential for axon development and brain wiring *in vivo* using *Drosophila melanogaster* as a model. *Drosophila* expresses a single ortholog of mammalian fascin called Singed (SN), which is expressed in the mushroom body (MB) of the central nervous system. Loss of SN causes severe MB disruption, marked by α- and β-lobe defects indicative of altered axonal guidance. SN-null flies also exhibit defective sensorimotor behaviors as assessed by the negative geotaxis assay. MB-specific expression of SN in SN-null flies rescues MB structure and sensorimotor deficits, indicating that SN functions autonomously in MB neurons. Together, our data from primary neuronal culture and *in vivo* models highlight a critical role for fascin in brain development and function.

**Highlights:**

- Fascin regulates axon growth and branching of hippocampal neurons in culture.
- Singed, a *Drosophila* fascin ortholog, is enriched in mushroom body (MB) axons.
- Singed loss causes axon guidance defects and sensorimotor issues in flies.
- MB-specific Singed re-expression rescues MB structure and behavior in flies.

## Introduction

The actin cytoskeleton powers a wide range of cellular activities ranging from cell migration to membrane trafficking^1,2^. The structure and dynamics of the actin cytoskeleton are regulated by over 100 actin binding proteins (ABPs)^3^. Several families of ABPs crosslink actin filaments (F-actin) to generate F-actin bundles of different densities that underlie distinct membrane protrusions such as microvilli and filopodia^4,5^. Filopodia are finger-like protrusions found on motile cells that play an important role in cell migration and cancer cell metastasis^6^. In developing neurons, they represent a hallmark feature of axonal growth cones, contributing significantly to axon guidance and branching^7–11^. Dynamic growth cone filopodia constantly survey the environment and respond to a complex array of external axon guidance signals, stabilizing when encountering attractive cues and retracting in response to repellent cues^12–15^. In support of a role for dynamic filopodia in axon guidance, eliminating growth cone filopodia was shown to abolish guidance responses *in vitro* and *in vivo*^16–18^. Additionally, filopodia are essential for collateral branching, a process vital for both neurodevelopment and injury recovery^9^. While the importance of filopodia in axon elongation, guidance, and branching is recognized, the molecular mechanisms underlying axon filopodia regulation and function remain to be fully elucidated.

While several families of ABPs can crosslink actin filaments to form different F-actin bundles, fascin is known to generate tight parallel F-actin bundles constituting the core F-actin structure in the filopodia of nerve growth cones and motile cells^19–22^. There are three mammalian isoforms of fascin, with fascin1 being the most widely expressed and studied^23^. Increased fascin1 expression in cancer cells is associated with increased metastasis and poor clinical prognosis, while knockdown of fascin1 leads to a decrease in cell migration^24–27^. Interestingly, fascin1 exhibits an intriguing expression profile in mammalian brains: it is highly expressed during development, but its expression is substantially decreased in the adult brain, suggesting an important role for fascin in early neurodevelopment^28,29^.

Despite its high level of expression in the developing brain and its importance in the formation and dynamics of filopodia in cell culture, the role of fascin1 in neural development and function, especially guided axonal development *in vivo*, remains unclear. An early study of gross brain morphology in a fascin1 knockout (KO) mouse model revealed an increase in the size of the lateral ventricles and the loss of the posterior region of the anterior commissure, suggesting a potential role for fascin1 in neuronal migration and axon development^30^. In addition, KO neurons exhibited smaller growth cones with fewer and shorter filopodia than wild-type control neurons in culture^30^. These findings, along with the high perinatal lethality exhibited by fascin1 KO mice (∼50%), suggest a crucial role in mouse development^30^. However, no detailed analysis of axon development, neuronal connections, synaptic function, or behavior in fascin1 KO mice was performed. Consistently, a separate fascin1 knockout mouse developed by the University of California, Davis Knock Out Mouse Project (KOMP) showed homozygous fascin1 KO animals exhibited a high degree of lethality^31^. Interestingly, the heterozygous mice were reported to exhibit neurobehavioral abnormality, further supporting a role for fascin in brain development and function^31^. In this study, we utilized a combination of mammalian cell culture and *Drosophila melanogaster* as an *in vivo* model to investigate the role of fascin in brain development and function. Our findings establish a critically important role for the fascin family of actin bundling proteins in brain development and function.

## Methods and Materials

### Cell culture

Dissociated primary hippocampal neuron cultures were prepared as previously described^32^. Briefly, 18.5-day old embryos were collected from timed-pregnant Sprague Dawley rats (Charles River Laboratories), and then hippocampi were dissected in ice-cold Hank’s Balanced Salt Solution (HBSS). Hippocampi were pooled together, trypsinized for 12 minutes (mins.), briefly incubated in 20% fetal bovine serum (Atlanta Biologicals, #S11150) and then plated on 25 mm coverslips coated with 100 μg/ml poly-L-lysine (P2636; Sigma-Aldrich) at two different densities, ∼150,000 or 25,000 cells per 35 mm dish for two different treatment conditions (see below). Hippocampal neurons were cultured in Neurobasal medium (21103049; Gibco) supplemented with 1% B-27 (17504044; Gibco), 1% Penicillin/Streptomycin (30002CI; ThermoFisher), and 1% GlutaMax (35050061; Gibco). The culture medium was replaced with fresh medium 24 hours (hrs.) after plating. Animal care and use was conducted following National Institutes of Health guidelines, and procedures were approved by the Institutional Animal Care and Use Committee at Emory University.

Cath.-a-differentiated (CAD) cells^33^ were used for analyses of fascin localization in lamellipodia and filopodia. CAD cells were cultured in DMEM/F12 (10-092-CV; Corning) media supplemented with 8% fetal bovine serum (Atlanta Biologicals, #S11150) and 1% Penicillin/Streptomycin (Invitrogen, #15140122). For imaging, we resuspended CAD cells by mechanical force and plated them on laminin-coated No. 1.5 glass coverslips for one hr. (laminin: CAS#: 114956-81-9, Millipore Sigma; 20 µg/mL for 1 hr. at 37 °C) before fixation and immunocytochemistry.

### Immunohistochemistry and microscopy of cultured cells

For immunofluorescent labeling of fascin and its cellular localization, both CAD cells and hippocampal neurons were fixed by cold (−20°C) anhydrate methanol (Sigma CAS# 67-56-1) for 10 min, followed by rehydration in 1x phosphate-buffered saline (PBS). Cells were blocked in PBS containing 2% normal donkey serum (D9663, Millipore-Sigma) for one hr. at room temperature. The cells were then incubated with primary antibodies at 4°C overnight and labeled with fluorescent secondary antibodies at room temperature for 1 hr. The following antibodies were used: fascin (1:1000 mouse anti-fascin, Sigma MAB3582, clone 55K2) and rabbit anti-β-actin (1:400, ab8227, Abcam). Alexa Fluor-conjugated secondary antibodies (Invitrogen) were diluted at a concentration of 1:500 in PBS-containing 2% donkey serum. All labeled coverslips were mounted on glass slides using Fluoromount-G (Southern Biotech #0100-01) for imaging.

Fluorescent imaging was mostly done using a Nikon Ti2 inverted microscope equipped with a Hamamatsu Flash4 scientific CMOS camera (for widefield) and a C2 confocal unit capable of simultaneous confocal imaging of three channels. Depending on the specific experiments, objectives of 10x/NA0.45, 20X/NA0.75, and 60X/NA1.4 (Plan Apo) were used. Structured illumination microscopy (SIM) was performed on a Nikon N-SIM system of Emory University Integrated Cellular Imaging Core Facility using a 100X/NA1.49 objective in 2D-SIM mode.

### NP-G2-044 treatment

The fascin inhibitor NP-G2-044 (HY-125506, MedChemExpress, Monmouth Junction, NJ) was stored at a concentration of 10 mM in DMSO. CAD cells were plated on 20 µg/ml laminin-coated 25 mm coverslips and allowed to attach and spread for one hr. CAD cells were then treated with either 15 µM of NP-G2-044 or 0.15% DMSO as a control. Coverslips were fixed at either 30 mins or 240 mins post-treatment application and subsequently labeled for fascin and β-actin. Images were acquired using widefield fluorescence microscopy. For this study, we selected fully flattened CAD cells with a “pancake” morphology to keep analysis consistent. For analysis, the segmented line tool in FIJI was used measure the average intensity of both the actin and fascin fluorescence in filopodia and the polygon tool was used to measure the cytosolic intensity of actin and fascin. The ratios of filopodial to cytosolic actin and fascin fluorescence intensities were calculated and analyzed via unpaired t-test for the means of the three biological replicates per condition.

To examine growth cone morphology, cultured hippocampal neurons (150,000/coverslip) were cultured for 3 days *in vitro* (DIV). The cells were then treated with 15 µM NP-G2-004 or 0.15% DMSO for 24 hrs., followed by methanol fixation and immunolabeling for fascin and β-actin on DIV4. Fluorescence microscopy and quantitative analyses were performed using the line tool in FIJI to trace the filopodia in each growth cone to determine length and number.

To investigate neurite extension, we cultured hippocampal neurons at a low density (∼25,000 cells/coverslip). On DIV1 the coverslips were inverted over a feeder layer of high-density rat cortical neurons (300,000/well) in 6 well plates with paraffin wax dots to separate the coverslips from the feeder layer. Starting DIV1, the neurons were treated with either 15 µM NP-G2-044, or DMSO vehicle control for 4 days, with fresh DMSO or NP-G2-044 added every day until fixation. On DIV5, the coverslips were removed, washed with PBS, fixed in 4% paraformaldehyde in PBS with 4% sucrose for 15 mins at room temperature, and then permeabilized in PBS containing 0.1% Triton X-100 for ten mins. The cells were blocked for one hr. at room temperature in 2% donkey serum (D9663, Millipore-Sigma) in PBS, and then incubated overnight at 4°C with mouse anti-tau (Sigma MAB3420, clone PC1C6) and rabbit anti-tubulin (Covance, PRB-435P) antibodies at 1:1000 in blocking buffer. Coverslips were next incubated in Alexa Fluor-conjugated secondary antibodies (Invitrogen A10037 and A11036) diluted in PBS 1:500 for an hr. at room temperature. Coverslips were then mounted in Fluoromount-G containing Dapi (Southern Biotech, Cat. No.: 0100-20). Axon length and the number of axon branch points were measured using the Simple Neurite Tracer (SNT) plugin for FIJI^34^. Slides were blinded for imaging and quantitative analyses. Statistical analysis was performed using GraphPad Prism Version 10.4.1 for macOS.

### Fly stocks

Fly stocks were obtained from Bloomington *Drosophila* Stock Center or were generously provided by Dr. Tina Tootle (University of Iowa), Dr, Kenneth Moberg (Emory University), and Dr. Jennifer Zanet (University of Toulouse, France). All crosses and stocks were maintained in standard conditions.

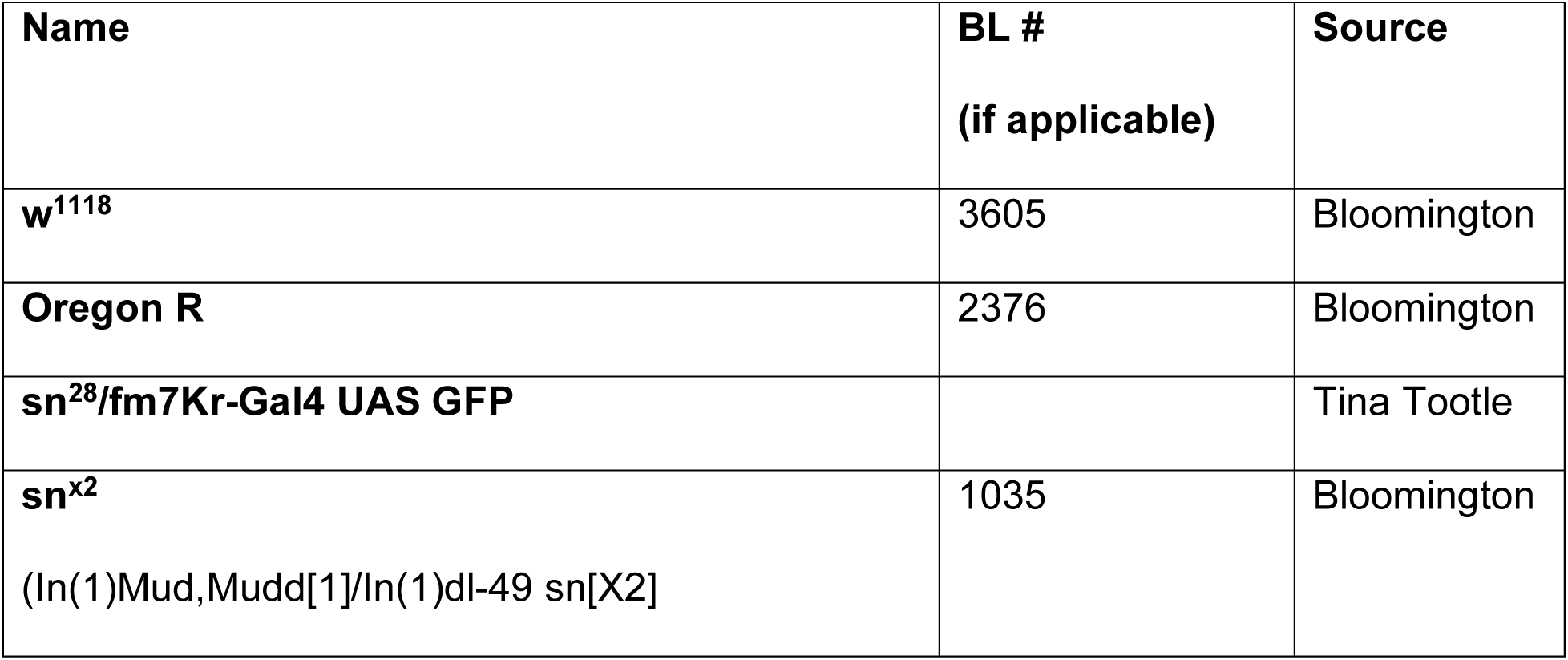
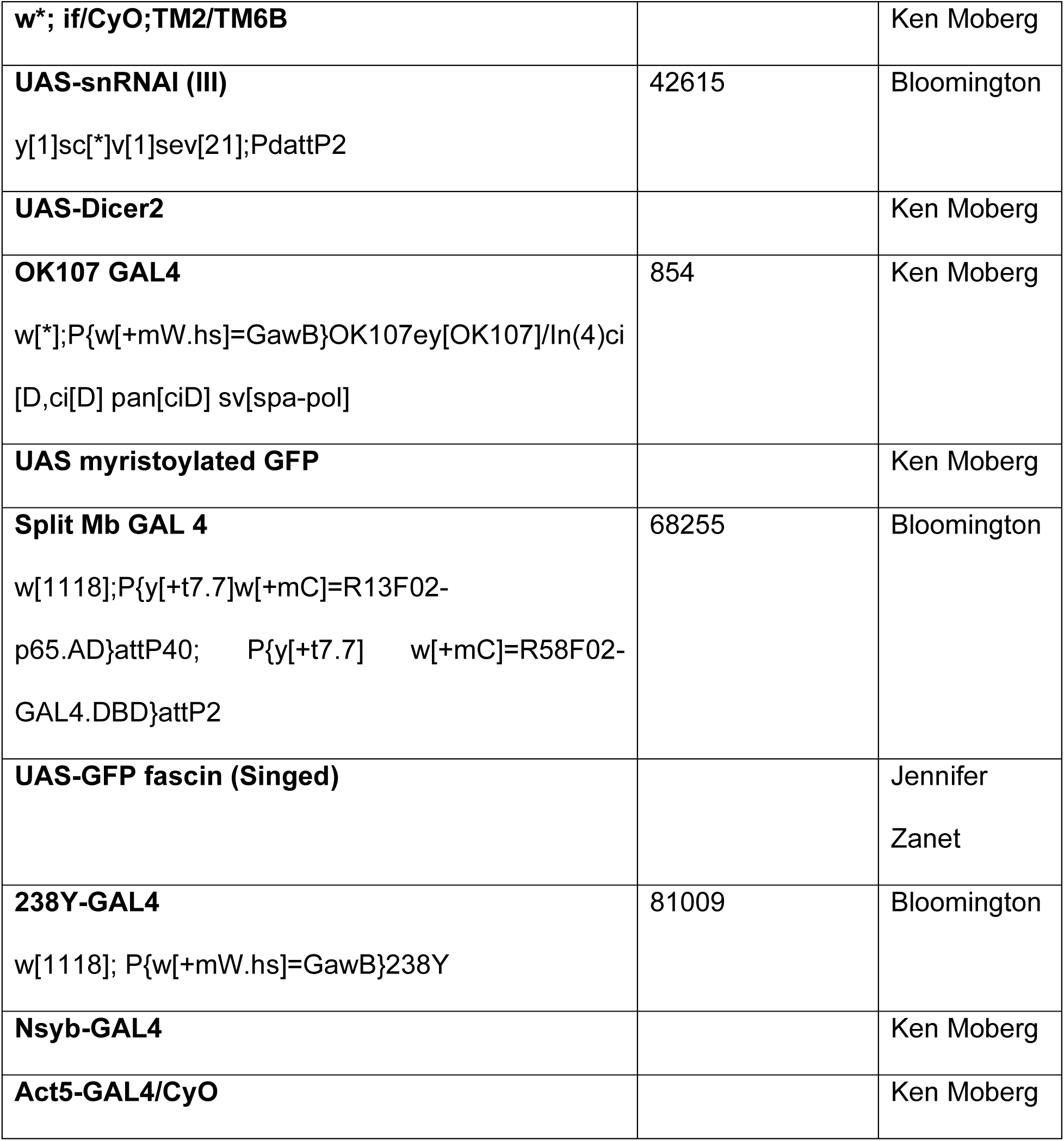

| Name | BL #<br>(if applicable) | Source |
| --- | --- | --- |
| **w<sup>1118</sup>** | 3605 | Bloomington |
| **Oregon R** | 2376 | Bloomington |
| **sn<sup>28</sup>/fm7Kr-Gal4 UAS GFP** |  | Tina Tootle |
| **sn<sup>x2</sup>**<br>(In(1)Mud,Mudd[1]/In(1)dl-49 sn[X2]) | 1035 | Bloomington |
| **w<sup>\*</sup>; if/CyO;TM2/TM6B** |  | Ken Moberg |
| **UAS-snRNAI (III)**<br>y[1]sc[*]v[1]sev[21];PdattP2 | 42615 | Bloomington |
| **UAS-Dicer2** |  | Ken Moberg |
| **OK107 GAL4**<br>w[*];P{w[+mW.hs]=GawB}OK107ey[OK107]/ln(4)ci<br>[D,ci[D] pan[ciD] sv[spa-pol] | 854 | Ken Moberg |
| **UAS myristoylated GFP** |  | Ken Moberg |
| **Split Mb GAL 4**<br>w[1118];P{y[+t7.7]w[+mC]=R13F02-<br>p65.AD}attP40; P{y[+t7.7] w[+mC]=R58F02-<br>GAL4.DBD}attP2 | 68255 | Bloomington |
| **UAS-GFP fascin (Singed)** |  | Jennifer<br>Zanet |
| **238Y-GAL4**<br>w[1118]; P{w[+mW.hs]=GawB}238Y | 81009 | Bloomington |
| **Nsyb-GAL4** |  | Ken Moberg |
| **Act5-GAL4/CyO** |  | Ken Moberg |

A balanced sn^28^ line was created by crossing sn28/fm7Kr-Gal4 UAS GFP flies with w*; if/CyO;TM2/TM6B flies. The resulting line was: sn28/fm7Kr-Gal4 UAS GFP; if/CyO;TM2/TM6B. This line was used to generate the Split-GAL4 and SN-rescue lines that are in the sn^28^ background.

The following genotypes were made through crosses and used experimentally:

1. Nsyb-GAL4; UAS myristoylated GFP
2. 238Y-GAL4; UAS myristoylated GFP
3. Split MB-GAL 4/ UAS myristoylated GFP; Split MB-GAL 4
4. sn^28^; Split MB-GAL 4/ UAS myristoylated GFP; Split MB-GAL 4
5. UAS-Dicer2/Act5-GAL4; UAS-snRNAi
6. UAS-Dicer2; UAS-snRNAi; OK107-GAL4
7. sn^28^; UAS-GFP Singed/238Y-GAL4
8. sn^28^; UAS myristoylated GFP/238Y-GAL4

### Fly brain dissection and staining

For anti-Fasciclin 2 (Fas II) and/or anti-GFP staining, dissections were performed as previously described in Behnke, *et al.* (2021)^35^. Briefly, whole flies were fixed for 3 hr. in 4% paraformaldehyde (PFA) in 0.01% Triton X-100-containing PBS (PBST). The flies were rinsed with PBST 4X for 15 min each, and brains were dissected in PBST. Brains were permeabilized overnight in 0.5% PBST at 4°C with nutation, then blocked in 5% normal donkey serum (D9663, Millipore-Sigma) in PBST for 90 min at room temperature with nutation. Brains were incubated in mouse anti-Fas II (DSHB, AB528235; 1:100) antibody and/or rabbit anti-GFP (Invitrogen, A11122; 1:1000) in blocking buffer overnight at 4°C with nutation. Brains were washed 3X with PBST and then incubated in secondary antibodies (Invitrogen A-11001 and A11032, 1:500) overnight at 4°C with nutation. The brains were then washed 3X with PBST and mounted in *SlowFade* Gold (ThermoFisher S36936) mounting media (ThermoFisher S36936).

Brains stained for SN were prepared using a protocol adapted from the FlyLight Double Label IHC for Adult *Drosophila* CNS^36^. Briefly, flies were anesthetized with CO_2_ and then put in PBS on ice. Brains were dissected in PBST and then stored on ice prior to fixation. The brains were fixed in 2% PFA in 0.5% PBST for 55 mins at room temperature with nutation. Brains were then washed 4x in 0.5% PBST for 10 mins and then blocked in 2% donkey serum (D9663, Millipore-Sigma) in 0.5% PBST for 90 mins at room temperature with nutation. Brains were incubated with anti-SN mouse antibody (DSHB, AB528239; undiluted) for 4 hrs. at room temperature with nutation and then incubated at 4°C with nutation for an additional 2 days. Brains were washed 4x for 15 min. in 0.5% PBST and then incubated in secondary antibody (Invitrogen A-11001 or A11032, 1:500) for 4 hrs. at room temperature followed by incubation at 4°C with nutation for 2 days. The brains were again washed 4x for 15 mins in 0.5% PBST and then mounted in *SlowFade* Gold mounting media.

### *Drosophila* Image Acquisition and Data Analysis

Unless otherwise noted, all data were collected from three independent biological replicates. *Drosophila* brains were imaged using an Olympus Fluoview FV1000-MPE two-photon confocal system with a Spectra-Physics MaiTi Deepsee Ti:sapphire femtosecond laser, using a 25X/1.05NA water immersion objective. Mushroom body defects were quantified by comparing blinded images to reference control images, as described previously in Kelly, *et al*. (2016)^37^. Representative control images were selected for male and female brains from either w^1118^ or Oregon-R flies based on previously reported characteristics of MB morphology. Images of brains from SN null and control flies were then blinded, compared to representative images, and scored for defects. The α- and β-lobes were scored in 4 categories: normal, thin, split (appearance of two axon distinct tracts), or missing. Midline defects were defined as present (either β-lobes crossing the midline or β-lobes failing to reach the midline) or absent (both β-lobes project to the midline and stop). GraphPad Prism Version 10.4.1 for macOS was used to calculate the N-1 Chi-squared test for comparison of two proportions for MB characterization, with each null line compared to its respective control line. A minimum of 25 brains were analyzed per genotype and 3 independent replicates were collected, except for the Split-GAL4 experiments. For comparisons between multiple proportions, Marascuilo’s Procedure was used post-hoc^38^.

For split-Gal4, a minimum of ten brains were analyzed per condition. N-1 Chi-squared test was used for statistical analysis for comparison of two proportions for MB characterization, with each null line compared to its respective control line.

### Modified Negative Geotaxis Assay (mNGA)

Data were collected as previously described in Ye, *et al.* (2024)^39^. Briefly, flies were placed in vials containing a layer of 2% agar at the bottom. Two to four vials were assayed at the same time using a custom 3-D Polylactic Acid (PLA) printed rig. In each trial, one vial contained control flies (either w^1118^ or Oregon-R) while the other contained SN null flies (either sn^28^ or sn^x2^) or SN null-rescue flies. Flies were separated and tested by sex. For each trial, the rig was lightly tamped three times, and fly movement was recorded using a Panasonic HC-V800 digital video recorder at 60 frames per second. Individual vials from each video were cropped, and the first 10 seconds were trimmed in Image J. Results were quantified by dividing the vial into 10 equal-size height bins and counting the number of flies in each bin every second after climbing begins. The climbing index was calculated based on the number of flies in each bin [see Ye, *et al.* (2024)^39^ for details]. All testing took place between ZT 3 and ZT 8 (ZT, Zeitgeber time; lights are turned on at ZT 0 and turned off at ZT 12) and testing occurred between 20 and 23°C under laboratory lighting conditions. The climbing index of each group of flies was plotted and statistically analyzed using paired T-tests for control versus null flies and two-way ANOVA with multiple comparisons for control, null, and rescue comparisons. Tracing images were obtained by creating Z-stacks of the video frames of the standard deviation of the fly positions five seconds into climbing.

### *Drosophila* western blots

Ten fly heads per genotype were collected and homogenized in RIPA buffer (150mM sodium chloride, 50 mM Tris-HCL, 1% Nonidet P-40, 0.5% sodium deoxycholate, 0.1% sodium dodecyl sulfate). The samples were spun down for 10 mins at 14,000 rpm, after which supernatant was collected. Samples were mixed 1:1 with 2x Laemmli buffer and run on a 4–20% Mini-PROTEAN® TGX™ Precast Protein Gel (BioRad, #4561094). After transfer to nitrocellulose, the membrane was blocked in 5% donkey serum in PBS + 0.2% Tween-20 for 1 hr. at room temperature. Blots were then incubated with the following primary antibodies in blocking buffer overnight at 4°: anti-SN mouse antibody (DSHB, AB528239; 1:50) and anti-α/β tubulin sheep antibody (Cytoskeleton, Inc. ATN02; 1:250). Blots were then incubated with the following secondary antibodies in blocking buffer for 1 hr. at room temperature: IRDye® 800CW Goat anti mouse IgG (Li-Cor, 926-32210; 1:1000) and Alexa Fluo3 647 donkey anti-Sheep IgG (Invitrogen, AB_2535865; 1:1000). Densitometry was performed using FIJI.

## Results

### Spatial distribution of fascin in polarized hippocampal neurons in culture

Fascin is the primary actin crosslinker in several membrane protrusions, including filopodia and microspikes, where it arranges up to 15-20 individual parallel actin filaments into tight bundles^19,40,41^. While previous studies have shown that fascin is enriched in filopodia to regulate their formation and dynamics^41,42^, recent studies suggest that fascin is also present in lamellipodia and may play a role in lamellipodial protrusions by regulating the organization and dynamics of branched F-actin networks^43–45^. To better understand the spatial distribution of fascin in motile membrane protrusions, we first performed immunofluorescent imaging of fascin distribution in mouse CAD cells in culture. CAD cells form large actin-rich membrane protrusions on laminin-coated surface, thus enabling high resolution examination of fascin association with distinct F-actin structures including those associated with lamellipodia and filopodia^46^. We found that fascin is enriched at filopodia and microspikes with thick F-actin bundles, as revealed by actin staining (Supplemental Figure S1, blue arrows). Importantly, we also observed a substantial level of fascin associated with branched F-actin networks in lamellipodia, especially at the leading edge (Supplemental Figure S1, red arrowheads). This observation is consistent with the notion that fascin may regulate branched F-actin networks underlying lamellipodial dynamics^43–45^.

The elaboration and extension of axonal projections during neural development depends on actin-based cell motility^10,12,47^. Over the first five days in culture, hippocampal neurons undergo stereotypical neurodevelopmental stages that result in a highly polarized morphology consisting of a single axonal projection and multiple minor dendritic processes^48^. Immunofluorescent imaging of fascin in hippocampal neurons on days in vitro 4 (DIV4) shows that fascin is highly enriched in the axon, particularly in axonal growth cones and at nascent axonal branches (Figure 1A&B, arrows). Consistent with previous reports in other neuronal cell types^20,21,49^, we found that fascin signals are concentrated in filopodia where β-actin is also enriched (Figure 1A color panels and Figure 1B, blue arrows). Noticeably, a substantial amount of fascin is observed in the lamellipodial region of the growth cone as revealed by confocal and super-resolution imaging (Figure 1A&B, red arrowheads), suggesting a potential role for fascin in the regulation of branched F-actin networks underlying growth cone lamellipodial protrusion. Consistently, exogenously expressed GFP-tagged fascin1 not only highlights growth cone filopodia but is also present in the lamellipodia (Supplemental movie S1).

**Figure 1.**
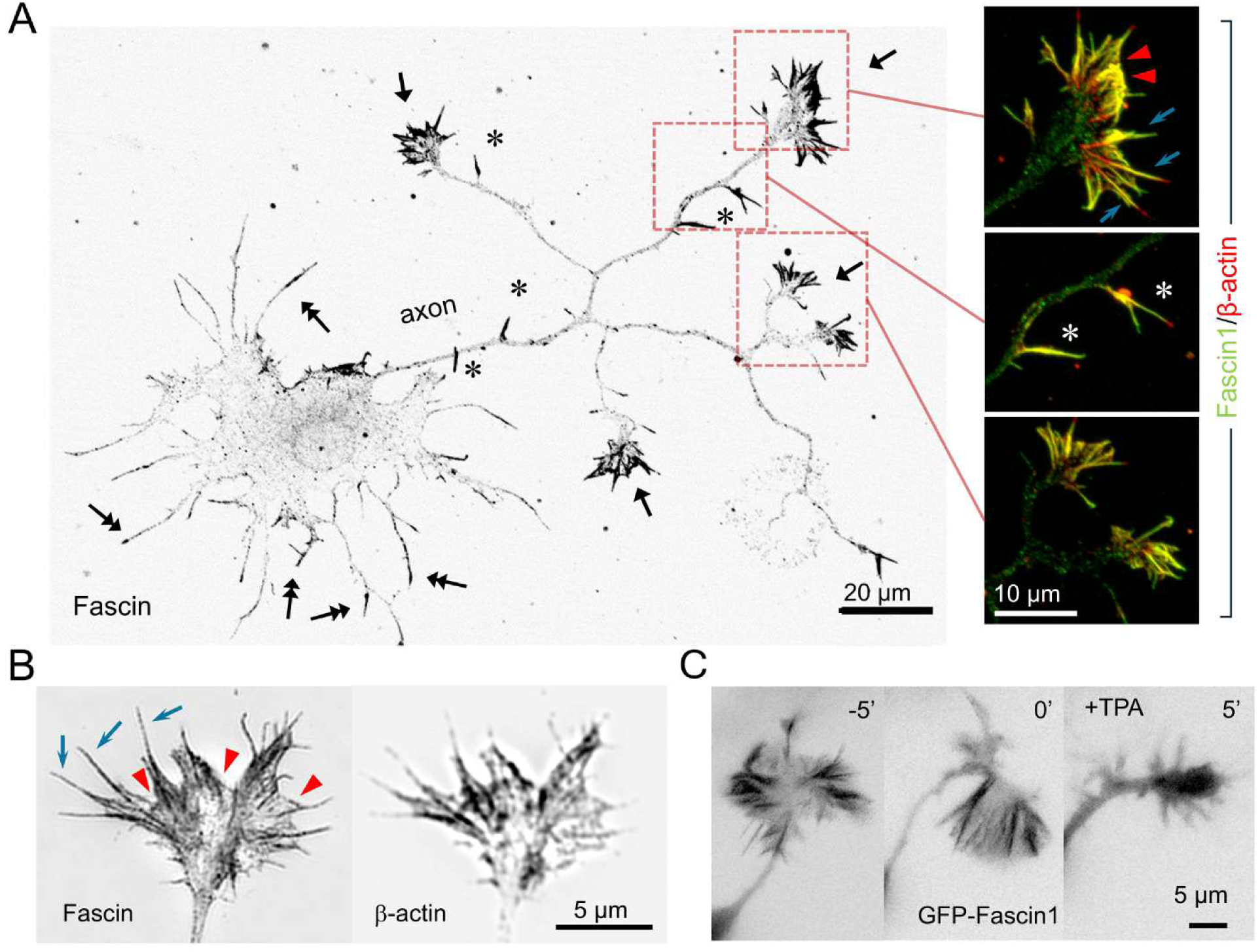
Fascin is concentrated in growth cone filopodia and collateral filopodia. (**A**) A confocal image of a DIV4 hippocampal neuron showing its polarized morphology with one axon and multiple minor processes. Neurons were methanol-fixed and stained for fascin and β-actin. A single axon with multiple branches has emerged from the soma. The black arrows indicate the enrichment of fascin in the axonal growth cones. Asterisks indicate the collateral filopodia along the axonal shaft. Two-headed arrows indicate the minor processes elaborated from the soma with substantial levels of fascin signals. The regions enclosed by dashed-line boxes are shown in a higher magnification in color with fascin in green and β-actin in red. Here, blue arrows indicate the fascin and β-actin enrichment in filopodia, and red arrowheads indicate the localization of fascin at the leading edge of the lamellipodia. (**B**) Structured illumination microscopy (SIM) images of a growth cone showing the presence of fascin in filopodia (blue arrows) and lamellipodia/leading edge (red arrowheads). (**C**) GFP-fascin localizes to growth cones and is mostly associated with filopodia. Five minutes after treatment with 12-O-tetradecanoylphorbol-13-acetate (TPA), a protein kinase C (PKC) activator, GFP-fascin is seen coming off filopodia, followed by filopodia retraction and resulting in growth cone collapse.

In addition to growth cones, filopodia protruding from the axonal shaft, referred to as collateral filopodia, are also enriched with fascin (Figure 1A&B, asterisks). Previous studies have shown that the formation of collateral filopodia along the axon is essential for branching^50^. Our observation does suggest a potential function for fascin in axonal branching. It should also be noted that many of the minor processes, which are believed to be the precursors of dendrites^48^, also contain a relatively high concentration of fascin (Figure 1A, two-headed arrows). While a previous study suggested that dendritic filopodia lack fascin^51^, it is plausible that fascin could be present and function during the early stages of dendritic development. Finally, consistent with previous reports^20,52,53^, activation of protein kinase C (PKC) by 12-O-tetradecanoylphorbol-13-acetate (TPA) causes the dissociation of fascin from F-actin bundles, leading to growth cone collapse (Figure 1C). This observation supports the notion that the cellular localization of fascin in the growth cones is associated with its F-actin bundling activity. Therefore, fascin likely plays an important role in actin-based axon development leading to the wiring of the brain.

### Inhibition of fascin’s actin bundling activity impairs growth cone filopodia, axon growth, and branching

To determine the role of fascin in growth cone motility and axon development, we utilized a potent small molecule inhibitor of fascin, NP-G2-044 (hereafter referred to as G2 for simplicity), which binds to fascin and prevents bundling activity^54,55^. We first tested the effects of this inhibitor in CAD cells by treating them with either 15 μM G2 or DMSO vehicle control for 30 mins. or 4 hrs., followed by methanol fixation and immunofluorescent staining to determine the effect of G2 treatment on the spatial distribution of fascin and β-actin. We observed that G2 treatment resulted in a gradual shift in fascin localization away from the actin-rich lamellipodia and filopodia at the cell edge and towards the central region of the cell (Figure 2A). We quantified the peripheral to central shift of fascin by measuring the ratio of the mean fascin fluorescence intensity at the cell edge over the mean fascin intensity in the center of the cell. At both the 30-minute and 4-hour timepoints the mean fluorescence intensity of fascin in the cell edge, relative to the cell center, significantly decreases in response to treatment with G2 (Figure 2B). This suggests that 15 μM G2 is sufficient to inhibit fascin-binding to F-actin in CAD cells. As a control, we also determined the ratio of β-actin intensity in the cell edge to the cytosol. We did not detect a shift in β-actin localization to the cytosol after treatment with G2 for the 30 minute timepoint but did observe a decrease in β-actin localization to the cell periphery for the 4 hour timepoint, indicating that the inhibitor affects fascin1 localization to the cell edge and that after 4 hours of treatment, the formation of F-actin structures at the cell edge may be decreased (Figure 2B).

**Figure 2.**
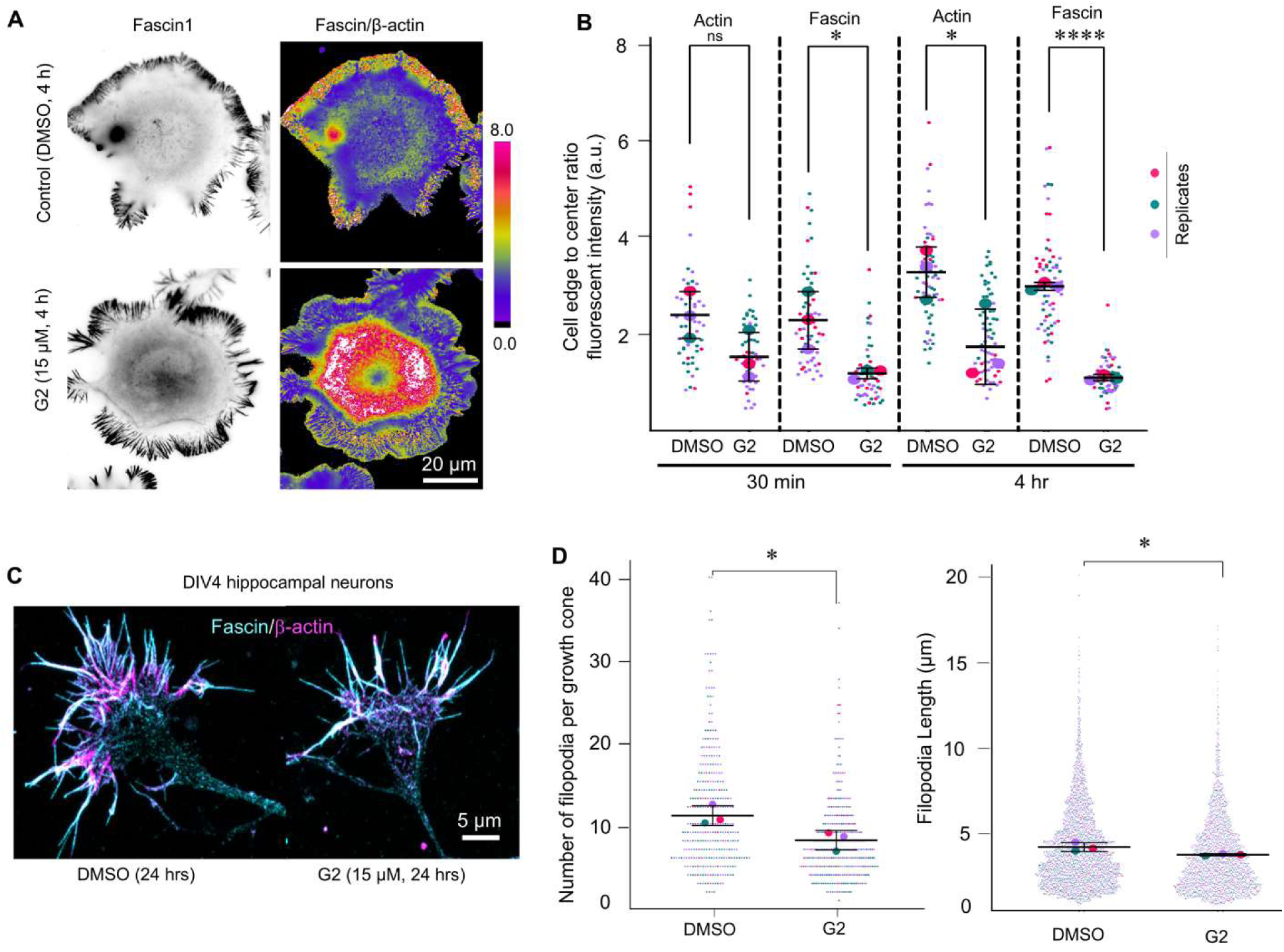
Inhibition of fascin-mediated actin bundling results in shift in localization and reduction in growth cone filopodia. (**A**) CAD cells were treated for 4 hrs. with either NP-G2-044 (G2) or DMSO and methanol-fixed. A decrease in fascin signal at the edge of the G2 treated cells and an increase in cytosolic signal is observed while β-actin signal does not undergo this shift in localization, as shown by ratio comparison. (**B**) Quantification of fascin and β-actin cell edge and center intensities shows a decrease in fascin localization to filopodia after application of G2 for both 30 minutes (p= 0.0340) and 4 hours (p<0.0001) when compared to cells treated with DMSO. β-actin signal does not undergo the same cytosolic shift after 30 minutes; however the 4-hour G2-treated neurons do show a significant difference in actin localization as well (p=0.0464). Statistical significance calculated via unpaired t-test for the means of the three biological replicates per condition, a minimum of 50 cells were analyzed per condition. Replicate one is shown in pink, replicate two in teal, and replicate 3 is in lilac. (**C**) Representative images of growth cone filopodia for DMSO control and G2-treated neurons. Fascin is shown in cyan and β-actin is shown in magenta. A decrease in number of filopodia as well as length of filopodia is seen. (**D**) Quantification of filopodia defects observed in fascin1-inhibited growth cones, and DMSO control growth cones (number of filopodia p=0.0372, filopodia length p=0.0421). Statistical significance calculated via unpaired t-test for the means of the three biological replicates per condition, with a minimum of 300 growth cones analyzed per condition.

We next investigated the effect of G2 treatment on filopodia morphology in neuronal growth cones by treating cultured rat hippocampal neurons on DIV3 with either DMSO or 15 µM G2 for 24 hrs., followed by methanol fixation and immunofluorescent labeling of fascin and β-actin on DIV4. After 24 hrs. of G2 exposure, axonal growth cones had significantly fewer filopodia (Figure 2C-D). In addition, the remaining filopodia were shorter in length (Figure 2C-D). These findings are consistent with what Yamakita, *et al.* (2009)^30^ reported in dorsal root ganglion neurons cultured from fascin1 KO mice, further supporting that G2 inhibits fascin activity in cultured neurons and led us to examine the effects of fascin-mediated actin bundling inhibition on axon outgrowth and branching.

We next applied G2 to DIV1 hippocampal neurons for four days to examine their axonal outgrowth and branching on DIV5. Here, neurons were treated with either 15 µM of G2 or DMSO as vehicle control. In mice, G2 was shown to have a half-life (T_1/2_) of ∼ 4 hrs.^56^ Therefore, to ensure that fascin was constantly inhibited for 4 days, fresh G2 or DMSO was applied to neurons every day. We fixed the neurons on DIV5 with PFA and stained for tau to visualize axons (Figure 3A), followed by tracing and quantification of axon length and branching for each condition (Figure 3B). We observed that neurons treated with 15 µM G2 exhibited reduced axon length and branching when compared to DMSO control neurons (Figure 3A-D). The reduction in length was mostly observed with secondary and tertiary branches, whereas the primary branch was slightly but not significantly affected (Figure 3C). Consistently, the number of secondary and tertiary branches was also significantly reduced in fascin inhibited neurons. Together with the localization of fascin to growth cones and collateral filopodia (see Figure 1), these data support the notion that fascin-mediated actin bundling promotes axonal growth and branching.

**Figure 3.**
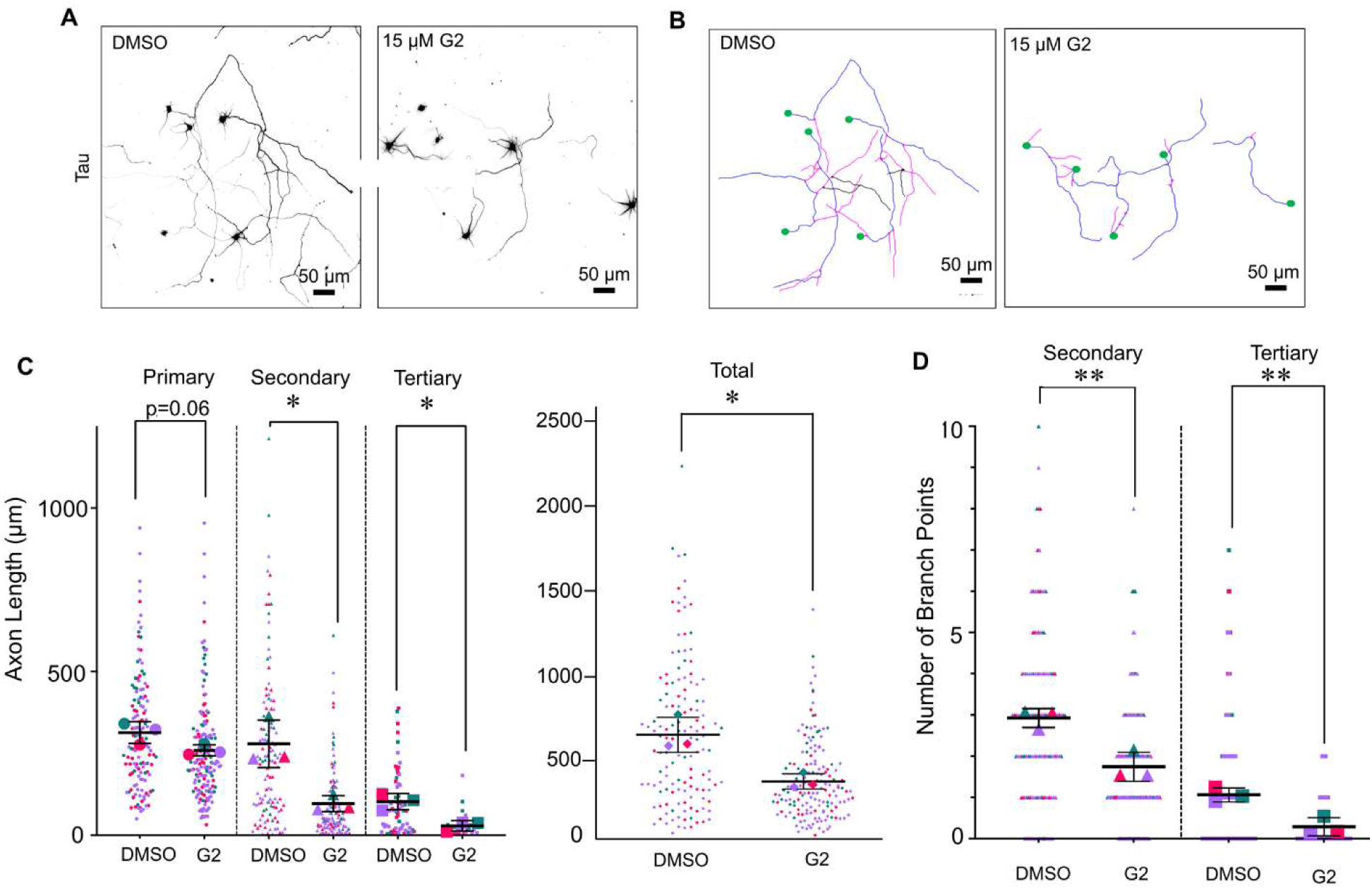
Pharmacological inhibition of fascin results in axon growth and branching defects. (**A**) Representative images of low-density primary rat hippocampal neurons PFA-fixed on DIV 5 and stained with anti-tau antibody. Neurons treated with 15 uM of G2 for 4 days show truncated axon growth and decreased branch points compared with DMSO-treated cells. (**B**) Tracings of representative DMSO control and G2-treated neurons, obtained via Simple Neurite Tracer for FIJI. Cell bodies are in green, primary axons are shown in blue, secondary in magenta, and tertiary in black. Neurons treated with 15 uM G2 show less axonal outgrowth and elaboration. (**C**) There are significant decreases in secondary (p=0.0145), tertiary (p=0.0125), and total (p=0.0134) axon length for the 15 uM G2-treated group compared to DMSO control neurons. Total secondary and tertiary axon lengths are shown. The primary axons for the fascin inhibited neurons showed a decrease that did not quite reach significance (p=0.0665). (**D**) Quantification of the number of secondary and tertiary branch points shows a significant decrease for neurons treated with 15 uM of G2 (p=0.0082 and p=0.0087, respectively). Statistical analyses were performed at the biological replicate level (n=3) using unpaired t-tests. Replicate one is shown in pink, replicate two in teal, and replicate 3 is in lilac.

### *Drosophila melanogaster* as an *in vivo* model to study axon development and brain wiring

Neurons in culture allow high-resolution imaging, easy manipulation, and quantitative assessment of their axonal growth and branching. However, they lack the complex and dynamic environment found *in vivo* that guides their axonal elongation to specific targets for precise brain wiring. To investigate how fascin regulates axonal development and brain wiring *in vivo*, we used *Drosophila melanogaster* as a model system. Unlike mammals, *Drosophila* expresses a single ortholog of fascin called Singed (SN), thus eliminating the possibility of compensation by other fascin isoforms. SN null flies exhibit defects in actin-based bristle morphology and female sterility due to defective actin structures in nurse cells, demonstrating that SN functions in F-actin bundling^57^. However, little is known about the expression pattern of SN in *Drosophila* brains and its potential functions in central nervous system development^57^. A previous study ^58^ showed that mushroom body (MB) γ-neurons isolated from SN null fly brains exhibit a filagree phenotype of neurite outgrowth in culture, suggesting a potential role for SN in axon development of MB γ-neurons. However, whether SN is expressed in MB neurons and other parts of the brain, as well as whether SN plays a role in MB development and wiring, remains completely unknown.

### SN is expressed in mushroom body neurons

Using an antibody against *Drosophila* SN, we detected SN in many neuronal cell bodies in the cortex and optic lobes of adult *Drosophila* brains (Figure 4A, arrows).

**Figure 4.**
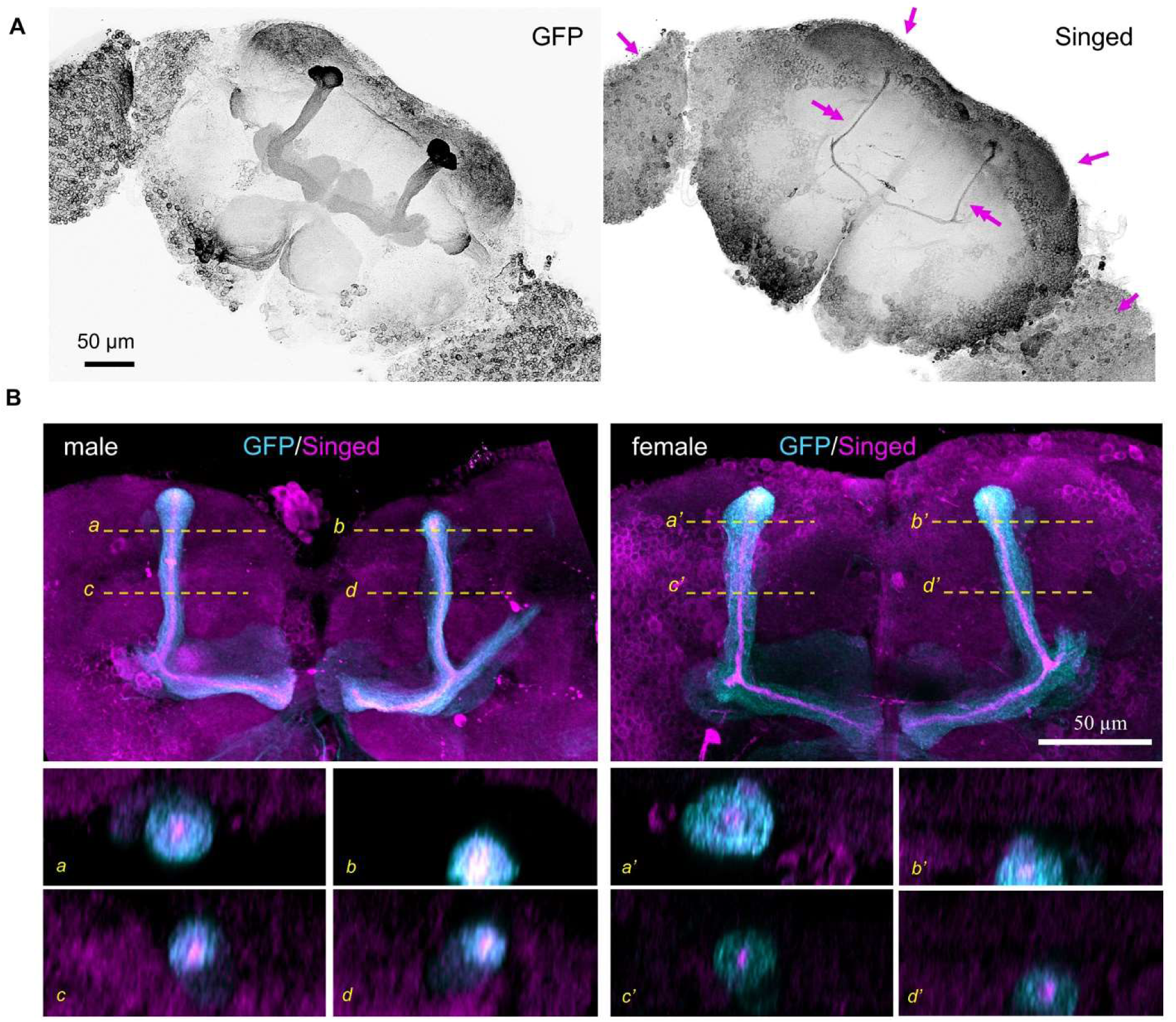
SN is expressed in a subset of MB ⍺- and β-lobes in adult *Drosophila* brains. (**A**) Brains from flies expressing pan-neuronal UAS-myristoylated GFP via the *nSyb-GAL4* driver were dissected and stained with anti-GFP and anti-SN antibodies. SN is seen in many neuronal bodies in cortex and optical lobes (arrows), as well as in the MB ⍺- and β-lobe axonal tracts (two-headed arrows). (**B**) Expression of UAS-myristoylated GFP via the 238Y GAL4 driver labels neurons in the MB born post-pupae formation and shows overlap with SN signal. GFP is labelled as cyan, and Singed is labelled as magenta. Axonal tracts labelled by SN staining appear to be centered in the MB axon bundles, as shown in the bottom cross sections. Yellow lines indicate the location of the cross section along the x-axis.

Strikingly, SN was found to highlight the mushroom body (MB) (Figure 4A, two-headed arrows), the part of the *Drosophila* central nervous system critical for olfactory learning, olfactory memory, and locomotion^59–61^. SN signals are enriched in both the α- and β-lobes, which are the axonal projections of Kenyon cells (KCs). Interestingly, the SN signal appears to be restricted within a narrow subset of MB axons. To confirm this, we used a MB-GAL4 (238-Y)^59^ line to drive expression of UAS-Myristoylated-GFP throughout the entire MB and then co-stained with anti-SN antibody. Our data revealed that SN is only expressed in axons found within the centers of the α- and β-lobes in adult brains (Figure 4B). Together, our findings confirm the expression of SN in the adult *Drosophila* brain and indicate the selective expression of SN in distinct populations of neurons not only within the central nervous system (CNS), but also within the MB in the adult *Drosophila* brain.

The development of MB starts in the early larval stages with γ-KCs elaborating their axons first to establish the initial framework of MB, which is followed by axons of α’/α- and β’/β-KCs that are born later^62^ (see also Supplemental Figure S2A). The narrow and centrally located SN-positive axons observed in α- and β-lobes of the adult brain appear to resemble the subset of KCs born later in CNS development that go on to form the cores of the α- and β-lobes^63^. We hypothesized that SN is more widely expressed in the *Drosophila* brain during early development, as it is in rodents and humans^28,29^. To test this, we fixed and stained 3^rd^ instar larval brains from flies expressing UAS-mCD8:GFP driven by the 201Y-GAL4^64^ line, which strongly marks MB γ-lobes in larval brains to examine SN expression (Supplemental Figure S2A). In comparison to the adult brain, SN is widely expressed in the cortex of larval brains, which overshadows the SN signals in MB when presented as maximal z-projection (Supplemental Figure S2B-C). In individual Z-sections through MB, SN signals can be clearly observed in γ-axons and KCs (Supplemental Figure 2D-E). In fact, SN and GFP signals (by 201Y GAL4-driven UAS-mCD8:GFP) appear to highly colocalized, indicating that SN is expressed essentially in all γ-axons during development. This observation supports the notion that the expression of Singed is more widespread during MB development than in the adult brain.

To investigate the function of fascin in *Drosophila* brain development, we obtained a null SN line (sn^28^) from Dr. Tina Tootle^65^. We first performed a western blot of female sn^28^ heterozygous and homozygous null fly brains to confirm SN knockout (Supplemental Figure S3A). We also confirmed via immunocytochemistry that SN is not expressed in sn^28^ brains (Supplemental Figure S3B). As a negative control we first looked at pigment dispersing factor (PDF) neurons, which do not appear to express SN (see Figure 4A), to see if there were any effects on their morphology. This subgroup of ventral lateral neurons belongs to the circadian clock neuron family and project axons from the optic lobe of the brain to the central brain^66^. As expected, we did not observe any differences in PDF neuron morphology between control and sn^28^ brains (Supplemental Figure 3C).

### Loss of SN results in axonal defects *in vivo*

The MB α- and β-lobes are formed by axons extended from KCs that undergo bifurcation and then project dorsally and medially (Figure 5A). The formation of MB lobes is dependent on both axon bifurcation and guidance, giving us a system to observe the effects of fascin depletion on both processes^67–71^. Brains were dissected from control w^1118^ flies and compared to sn^28^ flies. The brains were stained for Fas II, a marker for the MB α/β lobes^72^. We observed significant defects in the morphology of MB α- and β-lobes in SN null flies (Figure 5B). In control brains, the projecting axons bifurcate to form α- and β-lobes, with the α-lobes projecting dorsally and the β-lobes projecting medially and stopping at the midline of the brain. The control α- and β-lobes also appear to have tightly bundled axon tracks. In sn^28^ flies, we observe a range of defects, including missing, thin, or split (i.e. loosely bundled) α- and β-lobes (Figure 5C-D). We also observe a highly penetrative midline defect, with the β-lobes either crossing over the midline or in some cases failing to reach the midline (Figure 5E-F). The full statistical comparison of defects in control versus null brains is shown in Table 1. We conclude that global loss of SN results in significant MB α- and β-lobe defects.

**Figure 5.**
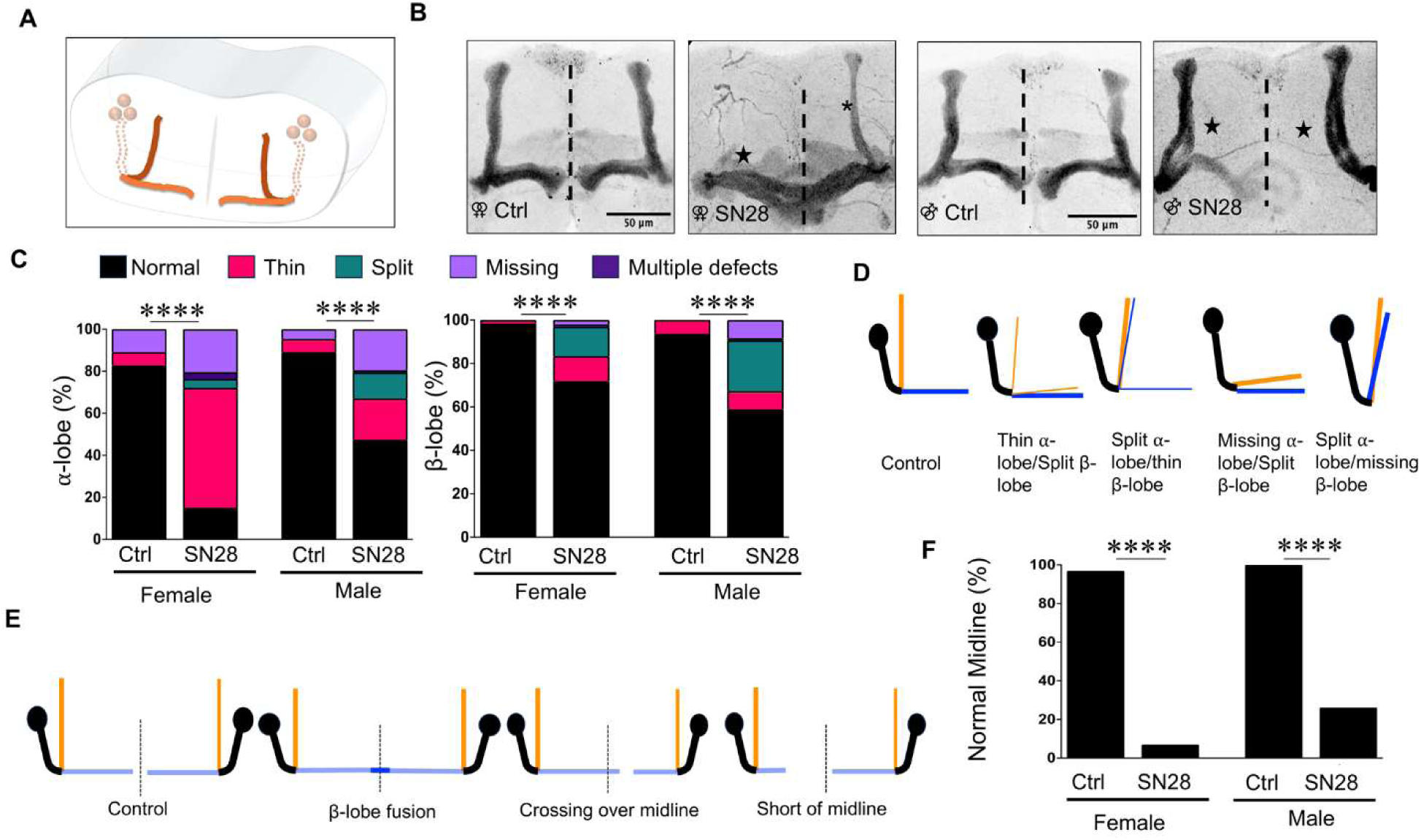
Loss of fascin results in axonal mushroom body (MB) defects. (**A**) Schematic of the Drosophila brain and the structure of the lobes making up the mushroom body (MB). The MB lobes are formed by projection of axons (dotted lines) from Kenyon cell bodies (spheres) which undergo bifurcation as well as guidance responses to form the dorsal ⍺- and medial β-lobes. (**B**) Brains were dissected from control (w^1118^) and SN null (sn^28^) flies and stained for Fas II, a marker of MB axons. The SN null flies display significant defects in both ⍺- and β lobes, as well as midline crossing errors (brain midline indicated by dashed black lines). Female w^1118^ brain, displaying normal ⍺- and β-lobes and no midline crossing of the β-lobes and female sn^28^ brain showing a missing ⍺-lobe, thin ⍺-lobe (black asterisk) and split β-lobe (black star). Male ^w1118^ and male sn^28^ brains, showing similar defects to the female brains. (**C**) Quantification of ⍺- and β-lobe defects. Chi square analysis was used to calculate p-values, shown in these graphs is the analysis of total defects (**** indicates a p-value of <0.0001). For a more detailed comparison of defects, see chart. (**D**) Schematics of ⍺- and β-lobe defects that we observe in SN null flies. (**E**) Schematics of the midline defects observed in SN null flies. **F)** Quantification of midline crossing defects in female w^1118^ (n=32), female sn^28^ (n=43), male w^1118^ (n=32), and male sn^28^ (n=42). Chi square analysis was used for statistical testing (p-value <0.0001).

**Table 1:**
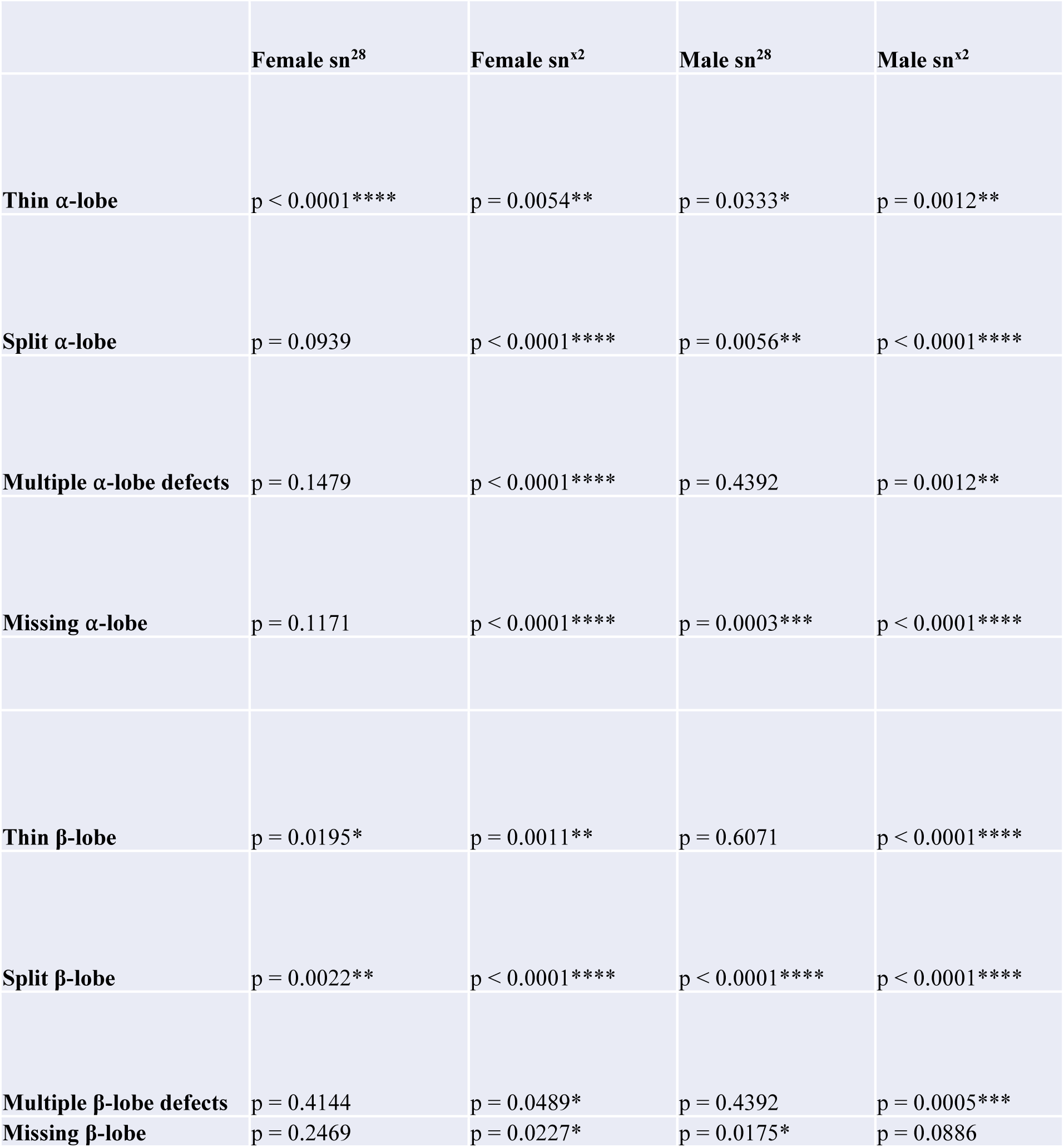
Statistical analysis of ⍺- and β-lobe defects in singed null flies. Chi square analysis was used to calculate p-values. sn^28^ flies were compared to control w^1118^ flies and sn^x2^ flies were compared to Oregon-R flies. Female w^1118^ n=32, female sn^28^ n=43, male w^1118^ n=32, and male sn^28^ n=42. Oregon-R n=54, female sn^x2 n^=43, male Oregon-R n=49, and male sn^x2^ n=34.

To better understand the MB defects in SN null fly brains, we employed a Split-GAL4 line to drive the expression of myristoylated GFP in a subset of MB axons^64^. In wildtype flies (Split-GAL4/UAS-myristoylated GFP; Split-GAL4/+), MB axons form stereotypical α- and β-lobes (Figure 6A, see Supplemental Movie 2 for 3-dimensional visualization). In sn^28^ flies (sn^28^; Split-GAL4/UAS-myristoylated GFP; Split-GAL4/+), we observed similar MB defects as described above, including MB lobes displaying thin, split, and missing phenotypes, as well as midline crossing defects (Figure 6B-C). Several of the guidance defects are highlighted by 3D rendering of the MB lobes in sn^28^ brains, including the midline crossing defects and split β-lobe (Figure 6B-C, orange and yellow arrows, see also Supp. Movie S3). We also observed several instances of β-lobe axons appearing to initially project medially, only to turn back towards the branch point (Figure 6C, white arrow, see also Supp. Movie S4) or project vertically (Figure 6C, orange arrow, see Supp. Movie S4). The MB lobe and midline defects observed in the sn^28^ flies expressing the split GAL4 driver are similar to those observed in the sn^28^ flies stained for Fas II (Figure 6D-E). These data indicate that SN is required for proper axon projection in the MB.

**Figure 6.**
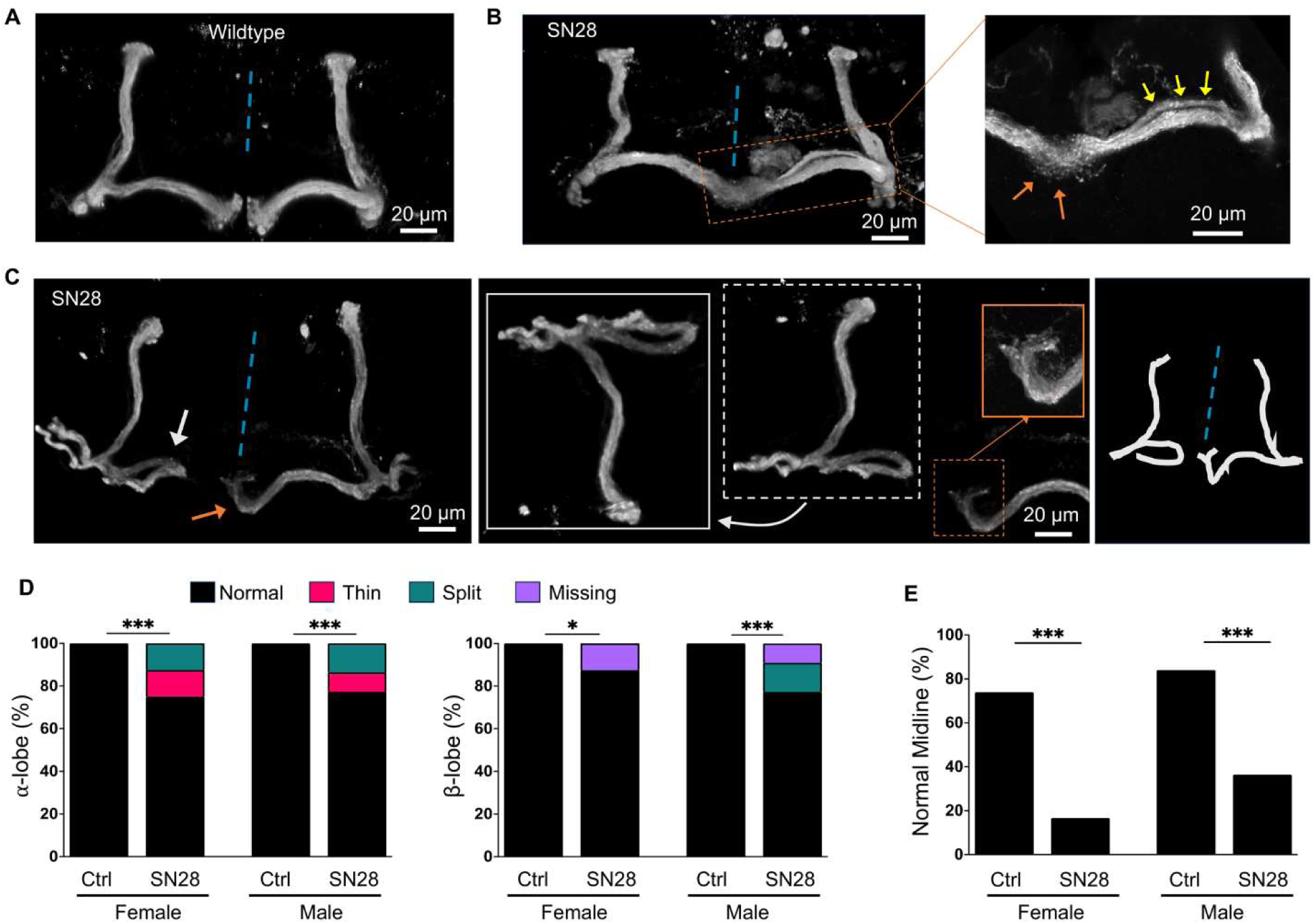
Fluorescent labeling of a subset of mushroom body axons reveals axon guidance deficiencies. Wildtype and sn^28^ flies expressing a MB-specific split GAL4 driver were crossed with wildtype or sn^28^ flies expressing UAS-Myristoylated GFP. Brains were dissected and stained for GFP and Fas II. (**A**) Wildtype MB axons project from Kenyon cells and undergo bifurcation to form two separate tracts (⍺- and β-) which then undergo axon guidance to form the vertical ⍺- and medial β-lobes. (**B**) sn^28^ MB axons undergo bifurcation but then fail to follow midline guidance signals, as shown by β-lobes crossing the midline (dotted blue line). To the right, the enlargement of the region outlined by the orange dotted box. We see the β-lobes crossing over the midline (orange arrows) and the split appearance of the β-lobe (yellow arrows). (**C**) Example of sn^28^ MB axons failing to properly follow guidance cues. White arrow shows a β-lobe turning back on itself, shown enlarged in panel on the right. The orange arrow points to a β-lobe that is starting to cross the midline (dashed blue line) and is also turning upwards at the midline, shown enlarged to the right. A schematic of the β-lobe trajectories is also shown. (**D**) Quantification of ⍺- and β-defects (female control n= 23, female sn^28^ n=12, male control n=25, male sn^28^=11), analyzed via Chi square analysis (p= 0.0004, p= 0.0007, p= 0.015, p=0.0007). (**E**) Quantification of midline defects, analyzed via Chi square (p= 0.0015, p=0.0048).

### Loss of SN results in sensorimotor defects

Previous studies have implicated the MB in locomotion with most MB output neurons having direct connections to the central complex, which controls locomotion and navigation in flies^60,73,74^. Consequently, locomotor defects are commonly associated with MB lobe defects^75,76^. We next used the modified negative geotaxis assay^39^ to assess if SN loss affects startle-induced climbing behavior. We found that there is a significant delay in climbing response for SN null flies when compared to controls, with null flies taking longer to reach the top of the vial than the control flies (Figure 7, Supp. Movie S5). Negative geotaxis represents a sensorimotor response that requires flies to first detect their position in respect to gravity and then move against the pull of gravity. Our data thus show that SN null flies exhibit significant sensorimotor defects as compared to control flies.

**Figure 7.**
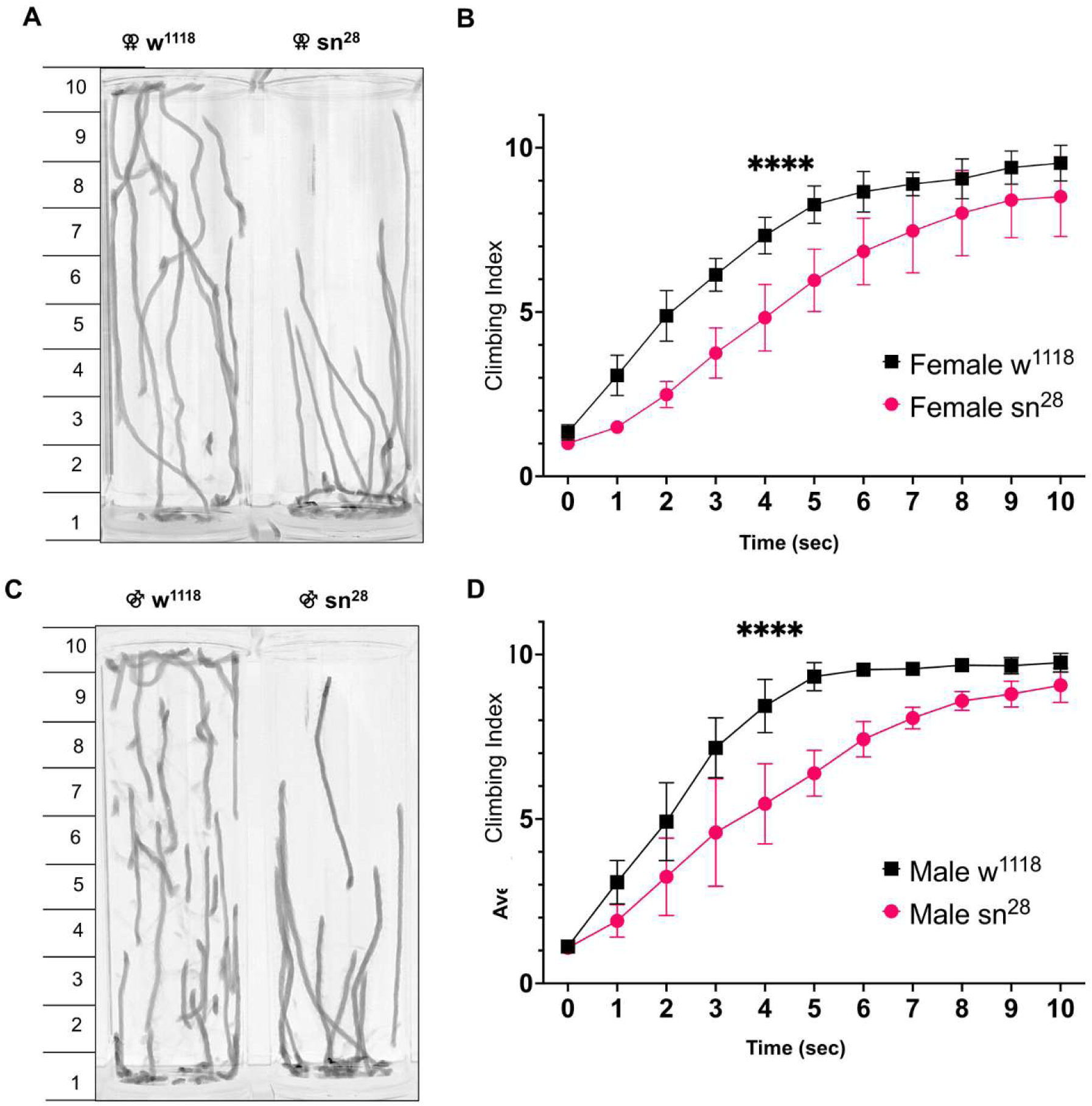
Singed null flies have a delayed climbing response. (**A**) Climbing behavior was evaluated using the negative geotaxis assay, flies are tamped to the bottom of a vial and results were quantified by dividing the vial into 10 bins (shown on the side of the image) and counting the number of flies in each bin 10 seconds after climbing begins Representative tracing of female w^1118^ and sn^28^ flies 5 seconds after beginning to climb. (**B**) Comparison of the climbing indices of w^1118^ (n=52) and sn^28^ (n=30) flies shows a climbing deficit for the sn^28^ flies, with the singed null flies taking longer to reach the top of the vial (analyzed using two-way ANOVA, p<0.0001). (**C**) and (**D**) w^1118^ male (n=48) and male sn^28^ (n= 33) flies show the same deficit seen in the female flies (analyzed using two-way ANOVA, p<0.0001).

To confirm the robust phenotypes observed in sn^28^ flies, we obtained a second SN null line (sn^x2^) to examine MB lobe morphology and the effects on sensorimotor behavior^57^. We used Oregon-R flies as the control line and found that sn^x2^ flies displayed similar defects in MB α- and β-lobe formation, midline crossing, and sensorimotor function (Supplemental Figure S4). Together, these data confirm the importance of SN for MB development and brain function.

### MB-specific depletion of SN results in α- and β-lobe defects

While SN appears to be strongly expressed in MB neurons and SN null lines display MB lobe defects, we wanted to determine if malformed MB lobes and deficits in startle-induced climbing were due to a MB neuron-specific requirement for SN or due to the global loss of SN in null flies. To answer this question, we knocked down SN expression specifically in MB neurons using the GAL4-UAS system to express an SN shRNA construct (snRNAi)^77^. For the initial characterization of SN knockdown, the Actin5-GAL4^78^ driver was used to globally express UAS-snRNAi and UAS-Dicer2^79^. We found that the expression of snRNAi in combination with Dicer2, resulted in a ∼60-70% reduction in SN protein levels as compared to control flies (Figure 8A). Flies that expressed both the UAS-snRNAi and UAS-Dicer2 constructs also displayed the gnarled bristle phenotype previously observed in SN null flies^57^ (Figure 8B). Next, we used OK107-GAL4^64^ to drive the expression of snRNAi specifically in MB neurons and their axons. We observed marked MB defects in UAS-Dicer2;UAS-snRNAi;OK107-GAL4 brains compared to control brains (Figure 8C-D), although they appear to be less severe than SN null brains. This is likely due to the incomplete KD of SN which is consistent with the lack of midline crossing defects in these brains. Our findings suggest that SN functions autonomously in MB to regulate its development and patterning.

**Figure 8.**
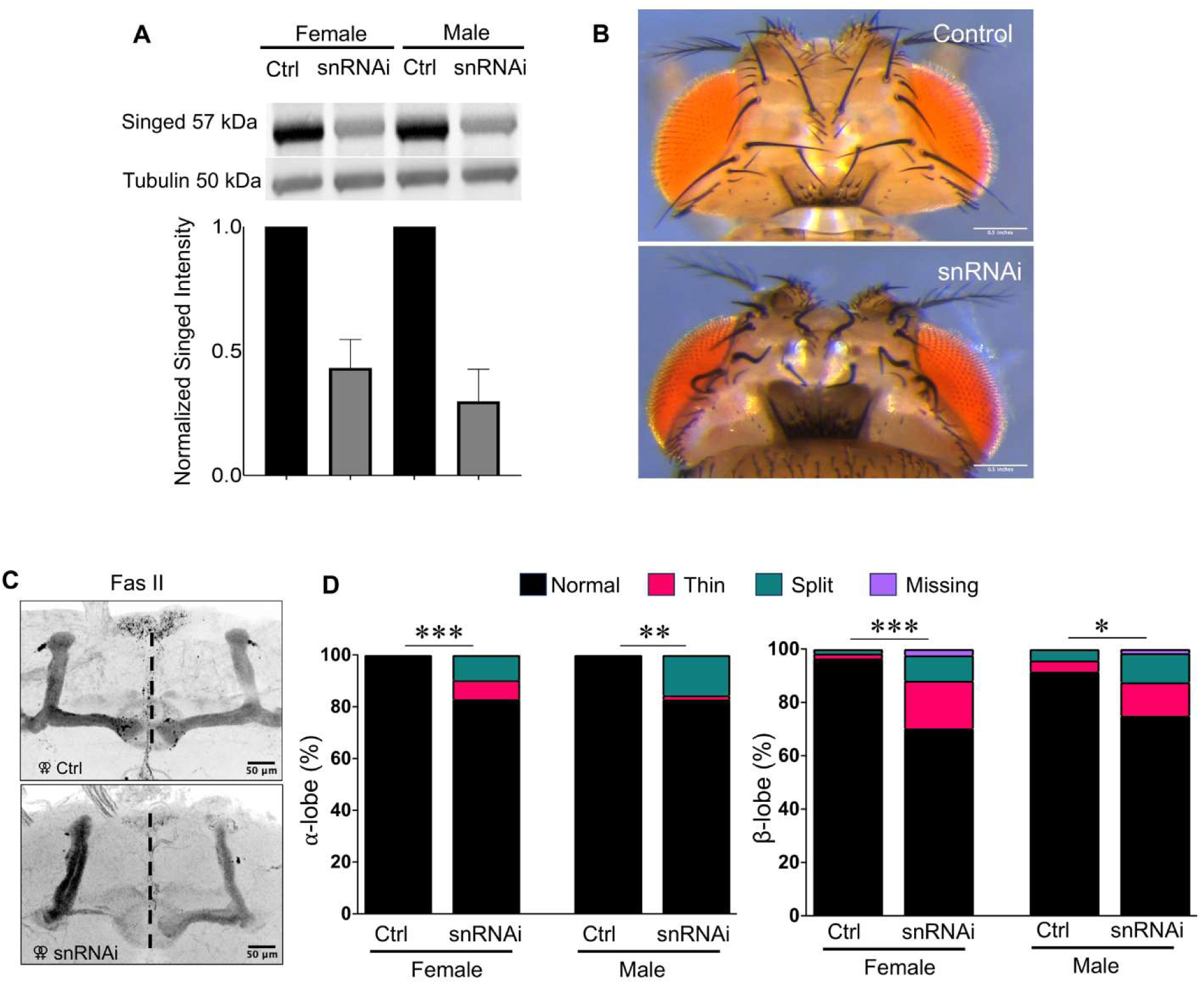
Mushroom body-specific knockdown of Fascin results in defects. (**A**) Quantification of Singed knockdown efficiency via western blot shows a decrease in SN expression but does not completely eliminate the presence of SN in heads collected from flies expressing a UAS-snRNAi construct as well as a UAS-Dicer2 construct ubiquitously expressed by an Actin GAL4 driver (n=3). (**B**) Flies expressing Act5-GAL4, UAS-Dicer2, and UAS-snRNAi display the gnarled bristle defect that is associated with loss of SN. (**C**) Representative images of control (UAS-Dicer2/CyO;UAS-snRNA/TM2) and snRNAi (UAS-Dicer2/+;UAS-snRNAi/+;OK107-GAL4 /+) brains stained for Fas II. (**D**) Quantification of ⍺- and β-lobe defects in female control (n=29), female MB KD (n=41), male control (n=25), and male MB KD (n=32). Midline (indicated by dashed black line) defects were not observed in the snRNAi brains. Chi square analysis was performed for comparisons, p values were as follows: p=0.0001, p =0.0261, p= 0.001, p= 0.003.

### MB-specific expression of SN is able to rescue defects

To further determine the requirement for SN in MB development, we used a MB-specific GAL4 driver (238Y)^59^ to drive the expression of either UAS-myristoylated-GFP (UAS myrs-GFP) as a negative control or wildtype UAS-GFP-Singed (UAS SN-GFP)^80^ constructs in the sn^28^ background. sn^28^ flies exhibiting the SN-null bristle defect^57^ and expressing both 238Y-GAL4 and either of the UAS-GFP constructs were collected and compared to control w^1118^ and sn^28^ flies in the mNGA and staining for MB morphology. We first examined startle-induced climbing in sn^28^ female flies expressing either UAS-myrs-GFP or wildtype UAS SN-GFP specifically in their MB neurons. Significantly, we found that MB-specific expression of SN-GFP in the sn^28^ background was able to restore startle-induced climbing to closely resemble control w^1118^ flies (Figure 9A). On the other hand, MB-specific expression of myrs-GFP in the sn^28^ background did not affect the impaired climbing behavior, and they exhibited the same deficits as sn^28^ flies (Figure 9A). Next, we examined whether MB-specific GFP-SN expression could rescue the structural MB defects in SN null flies. We found that the expression of UAS SN-GFP via the 238Y-Gal4 driver was sufficient to fully rescue MB α- and β-lobe defects to the levels seen in control w^1118^ flies (Figure 9B-C). Likewise, MB-specific SN-GFP expression successfully rescued the sn^28^ midline defects (Figure 9D). Together with our snRNAi results, these data indicate that SN functions in MB neurons to regulate the formation and patterning of α- and β-lobes during development.

**Figure 9.**
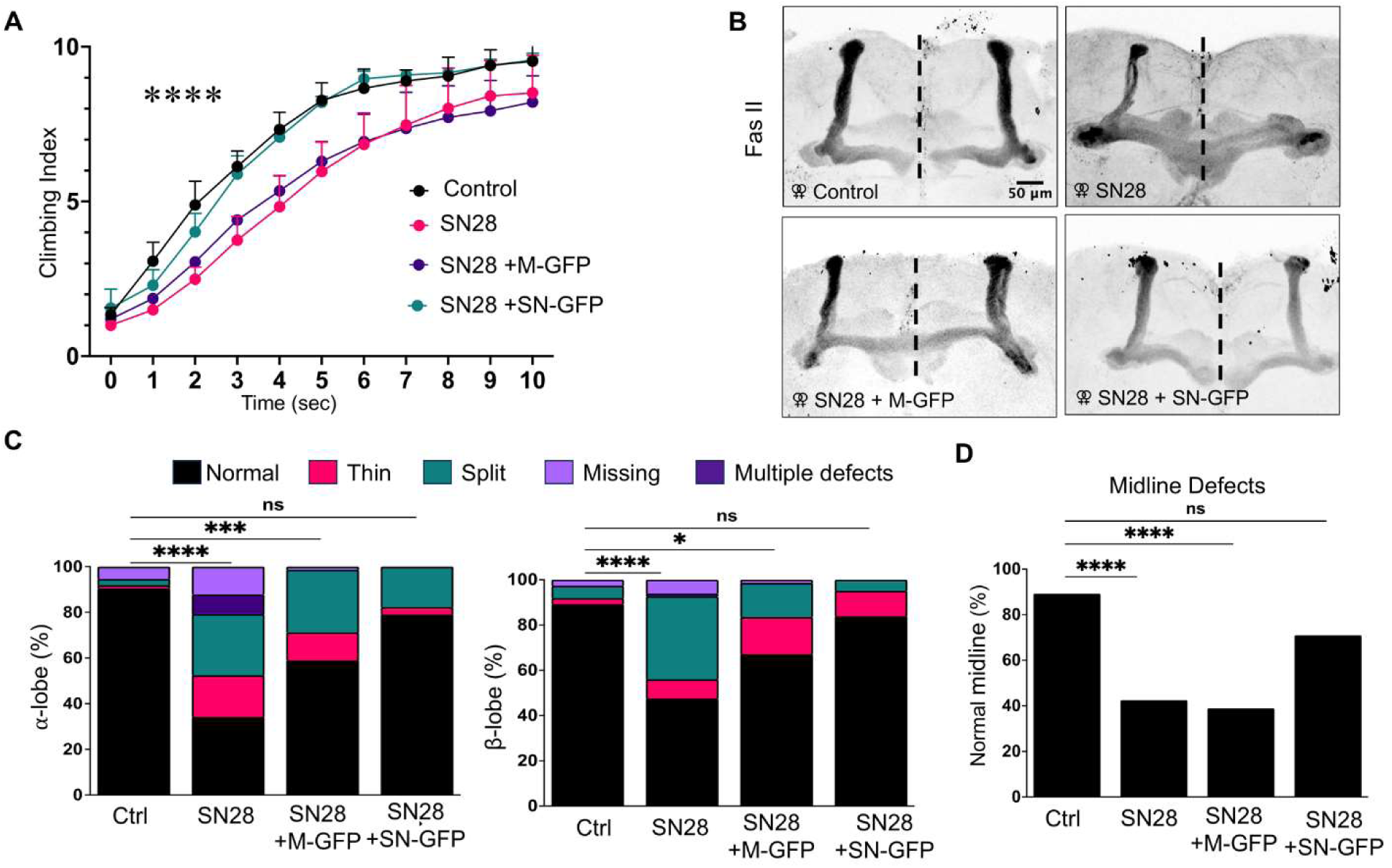
MB-specific expression of fascin1 is sufficient to rescue both climbing and MB lobe defects in females. (**A**) Female flies from 4 genotypes, w^1118^ (Control), sn^28^ (SN28), sn^28^;238Y-GAL4/UAS-Myristoylated GFP (SN28 + M-GFP), and sn^28^;238Y-GAL4/UAS-Singed GFP (SN + SN-GFP) were collected and assessed using the NGA protocol. Flies expressing UAS-GFP-Singed driven by 238Y-GAL4 are able to restore climbing ability to control w^1118^ levels (analyzed using two-way ANOVA, p<0.0001). (**B**) Representative images of MB ⍺- and β-lobes in the four genotypes examined. (**C**) Quantification of MB ⍺- and β-lobe defects shows that there is a complete rescue of MB lobe defects observed in sn^28^ flies upon MB-specific expression of WT GFP-Fascin1. Statistical analysis was performed using Chi-square proportional analysis with post-hoc Marascuilo’s Procedure, all experimental groups were compared to w^1118^ control. Female w^1118^ n=37, Female sn^28^ n=40 (β-lobe p-value <0.0001, ⍺-lobe p-value <0.0001), Female sn^28^; 238Y-GAL4/UAS Myristoylated-GFP n=36 (β-lobe p-value = 0.0465, ⍺-lobe p-value = 0.0004), female sn^28^; 238Y-GAL4/UAS Singed-GFP n=31 (β-lobe p-value = 0.9763, ⍺-lobe p-value = 0.6298). (**D**) MB-specific expression of UAS Singed-GFP is sufficient to restore midline axon projection morphology. Statistical analysis was performed using Chi-square proportional analysis with post-hoc Marascuilo’s Procedure. Female w^1118^ n=37, female sn^28^ n=40 (p-value <0.0001), female sn^28^; 238Y-GAL4/UAS Myristoylated-GFP n=36 (p-value <0.0001), female sn^28^; 238Y-GAL4/UAS Singed-GFP n=31 (p-value = 0.61).

It should be noted that MB-specific SN-GFP expression in male sn^28^ flies was also able to rescue both structural and behavioral deficits, but with reduced efficiency as compared to female flies (Supplemental Figure 5A-D). Whether there exists a sex difference in terms of SN-GFP expression or MB development remains to be investigated. Nonetheless, our *in vivo* data, together with results from cell culture, not only further demonstrate the importance of the fascin family of actin bundling proteins in actin based neuronal motility but also provide the first direct evidence that fascin’s role in developing neurons translates into higher-order functional outcomes.

## Discussion

As a major F-actin bundling protein, fascin is concentrated in protrusive cellular structures such as invadopodia and filopodia that play critical roles in directed cell motility and invasion of the cellular matrix^12,19,20,23,81–83^. Previous studies in culture have established that fascin is concentrated in growth cone filopodia and regulates filopodia dynamics^2,20,21^. Given that growth cone filopodia are considered the sensory motile apparatus for guided axonal growth, it is conceivable that fascin plays a crucial role in axon development and brain wiring^11,13^. However, direct evidence to support this conclusion remains lacking. In culture, an increase in fascin in the growth cone of mouse dorsal root ganglion (DRG) neurons was observed in response to the repulsive cue Semaphorin 3A, but whether this change in the fascin level is required for growth cone collapse was not investigated^49^. *In vivo*, a fascin1 knockout (KO) mouse model revealed grossly normal brain structures with the exception of an increase in the size of the lateral ventricles and the loss of the posterior region of the anterior commissure, suggesting a potential role for fascin1 in neuronal migration and axon development^30^. Explant DRG cultures from the fascin KO mouse were able to extend neurites with similar lengths to that of the wild type but exhibited smaller growth cones with fewer and shorter filopodia^30^. While fascin1 KO has been shown to disrupt neuronal migration^84^, whether fascin is required for guided axonal projection, axonal branching, and brain wiring was undetermined.

In this study, we take advantage of mammalian primary neuronal cultures and a *Drosophila* model, combined with pharmacological, genetic and imaging approaches, to establish that fascin plays a critical role in axonal development and brain wiring. Here, primary mammalian neurons in culture have enabled us to examine the distinct cellular localization of fascin in developing neurons, and to understand how fascin-mediated actin-bundling regulates axon outgrowth and branching. Additionally, the use of *Drosophila* as an *in vivo* model reveals the critical importance of fascin in axon development, brain wiring and function *in vivo*. Through a combination of genetic manipulation, whole brain imaging, and sensorimotor behavioral assays, we provide direct evidence that fascin is required for the proper development of the mushroom body (MB), which is analogous to the mammalian hippocampus^85^. Loss of fascin in *Drosophila* leads to defects in both α- and β-lobes, which represent guided axonal projections from Kenyon cells^72^. We additionally demonstrate that the loss of fascin leads to deficits in startle-induced climbing. Importantly, MB-specific expression of fascin is sufficient to rescue observed structural and behavioral deficits. Together, our study has established a critically important role for fascin-mediated actin bundling in brain development and function.

### Fascin in developing neurons

Primary rat hippocampal neurons in culture represent a well-established model to investigate axon formation as they undergo distinct stages leading to the establishment of a single axon within a one-week period^48^. Importantly, hippocampal neurons in culture allow high-resolution imaging that not only helps to determine the cellular localization of fascin at various stages of axon development but also enables the tracking and quantification of axonal branching and growth of individual neurons. In cultured hippocampal neurons, we found that fascin is highly enriched in growth cones, especially growth cone filopodia. Importantly, fascin signals were also detected in growth cone lamellipodia, suggesting a potential role in the regulation of branched F-actin networks underlying lamellipodial protrusions^43–45^. It should be noted that pharmacological elimination of filopodia was previously shown to abolish directed guidance responses with a slight reduction in neurite elongation^17,18^. Therefore, our finding that fascin inhibition reduced the axonal length supports a potential role for fascin in regulating filopodial actin dynamics underlying axon extension.

Our imaging also revealed fascin enrichment in collateral filopodial protrusions, suggesting a potential role for fascin-mediated actin dynamics in axonal branching. Axonal branching is an important developmental event that enables a neuron to establish synaptic connections with multiple targets^86^. Axon branching occurs through two main mechanisms. The first is when growth cones split to form two separate axon paths, also known as bifurcation, and the second is collateral branch formation, which occurs when new branch points are established on the axon shaft^87^. For either mechanism, the presence of filopodia is important to initiate branch formation^88^. Indeed, inhibition of fascin by the small molecule inhibitor NP-G2-044 prominently decreased axon branch points and the outgrowth of secondary and tertiary axon branches, supporting a role for fascin in axon branching and growth. NP-G2-044 is a well-established and highly-specific fascin inhibitor^54^. Importantly, crystal structure and cryo-electron microscopy studies have provided a mechanistic understanding of how and where this small compound binds to fascin to interfere with its bundling activities^89,55^. As a result, the rapid action of fascin inhibition by NP-G2-044 has enabled us to examine the immediate, short-term, and long-term effects on axonal development in culture. The subsequent use of a *Drosophila* model combined with genetic manipulation has substantiated our work on cultured neurons and allowed us to elucidate the *in vivo* function of fascin.

### Expression and roles of fascin in the mushroom body

In *Drosophila,* SN is the only ortholog of fascin and is required for actin bundles in bristles^90^, nurse cells^57^ in female flies, and dendritic terminal branching of class III sensory DA neurons^91^. However, the expression pattern of SN in *Drosophila* brains and its function in brain development were not well studied. Previous work^58^ found that loss of SN resulted in filagree shaped neurites from cultured MB γ-neurons, suggesting that SN may function in axonal development, at least *in vitro*. However, whether SN is expressed in the MB or how it affects MB development was not investigated. We have shown for the first time that SN is expressed in a narrow subset of MB α- and β-lobe axons in the adult *Drosophila* brain, which we suspect to represent the axons of late-born neurons^63^. The development of MB in *Drosophila* mostly occurs in larval and pupal stages, with γ-axons leading the way^62^. Consistent with a role for fascin/SN in axon development, we observed widely expressed SN in larval brains and importantly in almost all γ-axons highlighted by 201Y GAL4-driven UAS-mCD8:GFP. This observation is consistent with the high expression profile of fascin during brain development in rodents and humans^28,29^. Given that γ-axons appear to serve as pioneer axons^62^, loss of SN in these neurons could lead to the disruption of α- and β-lobe formation. Future work could utilize different GAL4 drivers to deplete SN in specific subsets of developing neurons to determine if SN presence is required in all MB neurons for proper lobe formation.

The observed defects in MB lobes most closely resemble guidance errors. MB axons are guided by multiple guidance factors and signaling pathways^67–69,92–94^. One of the most well-studied and conserved pathways is the Slit/Robo family which governs midline crossing in *Drosophila*^94^. Previous work in mammalian cultured neurons has shown that Slit-mediated repulsion is dependent on filopodia dynamics^95^. Here, we show that loss of fascin in MB axons results in a highly penetrative midline defect. This further supports a role for fascin’s regulation of filopodia in axon guidance. Our *Drosophila* Split-GAL4 data show that the axon paths extending from Kenyon cell bodies appear to successfully bifurcate and then fail to follow proper guidance pathways, hence the prominence of the “split lobe” phenotype that we observe in SN null brains. We speculate that this phenotype is due to α-lobe axons following β-lobe guidance pathways and vice versa. The determination of which axons become α-lobe and which become β-lobe is dependent on tightly regulated expression and localization of the Eph/Ephrin family of guidance molecules^70^. The interaction of fascin and this family has not been studied in neurons, however a study in prostate cancer cells showed that the addition of an ephrin ligand resulted in an increase in fascin-positive filopodia^96^, suggesting that the split lobe phenotype we observed could be due to disruption in response to Eph/Ephrin signaling. At this moment, we are unable to definitively determine if MB axon bifurcation is affected by SN loss. Future experiments at the single cell level, such as mosaic analysis with a repressible cell marker (MARCM)^97^, will likely provide the answer.

### Fascin’s function in axon development and brain wiring

Developing axons are guided by a myriad of attractive and repulsive cues that are surface-bound or diffusible in nature^94,98–106^. It is believed that the motile growth cones at the tip of axons sense the environment via dynamic filopodia and steer the axons through actin-based motility^16–18^. Given that fascin is enriched in growth cone filopodia, it may serve as an important cytoskeletal component to support the formation and extension of filopodia. On the other hand, fascin can be phosphorylated at serine 39 by protein kinase C (PKC), and S39 phosphorylation diminishes its bundling activity^52,53^. An additional phosphorylation site of fascin (S274)^107^ has also been identified along with two lysines (K247, K250) that can be monoubiquitinated^108^. Therefore, fascin-mediated F-actin bundles underlying filopodial protrusions can be dynamically regulated in response to signaling cascades during axon pathfinding. Furthermore, fascin has been shown to interact with Dishevelled-associated activator of morphogenesis (DAAM1), a member of the formin family^109^. DAAM functions as a link between the Wnt5/planar cell polarity (PCP) guidance system and the actin cytoskeleton, which guides MB lobe projection^69^ in *Drosophila* neurons. DAAM1 is also enriched in the newly formed axons of MB neurons, where it is required for proper lobe projection^110^. Finally, both DAAM1 and fascin1 have been shown to bind to microtubules and actin,^111,112^ raising the possibility that fascin may coordinate these two cytoskeletal systems for guided axon elongation during brain wiring. Therefore, it is reasonable to postulate that fascin may function beyond making F-actin bundles for filopodia and be actively involved in the intricate signal transduction that underlies axon guidance and brain wiring.

## Supplemental materials

### 1. Supplemental methods

#### Expression of GFP-fascin1 in neurons

Rat hippocampal neurons were cultured as described in the methods section of this paper. On DIV4, they were transfected via the calcium phosphate method^113^ with 3 ug of a GFP-fascin1 construct generously given by Danijela Vignjevic^19^. Widefield fluorescent time-lapse imaging of GFP-fascin1 was performed 24 hours after transfection (on DIV5) using the Nikon Ti2 system.

#### Larval brain staining

Wandering third instar larvae from y[1] w[67c23]; P{w[+mC]=UAS-mCD8::GFP.L}LL5 P{w[+mW.hs]=GawB}Tab2[201Y] (Bloomington Drosophila Stock Center, RRID:BDSC_64296) flies were dissected in phosphate-buffered saline containing 0.01% Triton-X (PBST). Brains were then fixed for 40 mins. in 2% paraformaldehyde (PFA) at room temperature with rocking. After fixation, the brains were washed 3X in PBST and then blocked in PBST + 2% donkey serum overnight at 4°C with nutation. Primary antibodies were incubated for 4 hrs. at room temperature followed by two days at 4°C with nutation. The following primary antibodies were used: anti-SN mouse antibody (DSHB, AB528239; undiluted) and rabbit anti-GFP (Invitrogen, A11122; 1:1000). The brains were washed 3X in PBST and incubated with secondary antibodies for 4 hrs. at room temperature followed by 2 days 4°C with nutation. The following secondaries were used: Invitrogen A-11001 and A11032, both at 1:500. The brains were then washed 3X in PBST followed by 3X washes in PBS and finally mounted in *SlowFade* Gold mounting media (ThermoFisher S36936).

#### PDF neuron staining

Whole flies were fixed for 3 hrs. in 4% paraformaldehyde PFA in PBST. The flies were rinsed 4 times for 15 mins. each with PBST and brains were subsequently dissected in PBST. Brains were permeabilized overnight in 0.5% PBST at 4°C with nutation then blocked in 5% normal goat serum in PBST for 90 mins. at room temperature with nutation. After blocking, the brains were incubated in mouse anti-PDF (DSHB, PDF C7, AB_760350 AB_2315084) antibody diluted 1:100 in blocking buffer overnight at 4°C with nutation. Brains were washed 3 times with PBST and then incubated in secondary antibody (Invitrogen A-11001) overnight at 4°C with nutation. The brains were then washed 3 times in PBS and mounted in *SlowFade* Gold mounting media (ThermoFisher S36936).

### 2. Supplemental figures (S1-S5)

**Figure S1:**
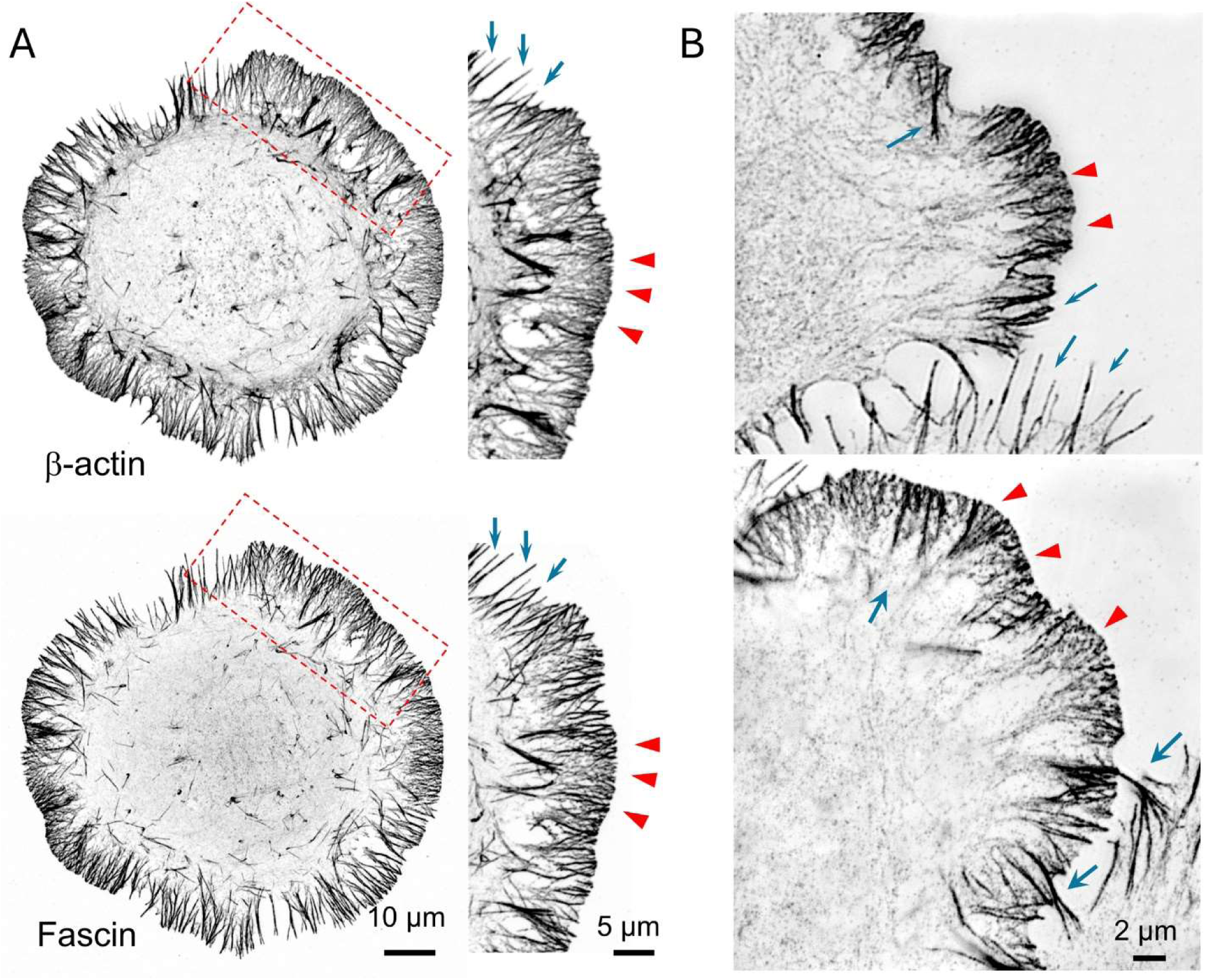
Fascin enrichment in filopodia and lamellipodia and its association with different F-actin structures in CAD cells. (**A**) Confocal images of a CAD cell stained β-actin (top row) and fascin (bottom row). Regions enclosed by red dashed-line rectangles are enlarged and shown on the right. Fascin is clearly associated with thick F-actin bundles such as filopodia and microspikes (blue arrows). However, a substantial level of fascin signals is also seen in the lamellipodia (red arrowheads). (**B**) Structured illumination microscopy (SIM) further reveals the fascin localization to thick F-actin bundles (blue arrows) and the F-actin meshwork in lamellipodia (red arrowheads).

**Figure S2:**
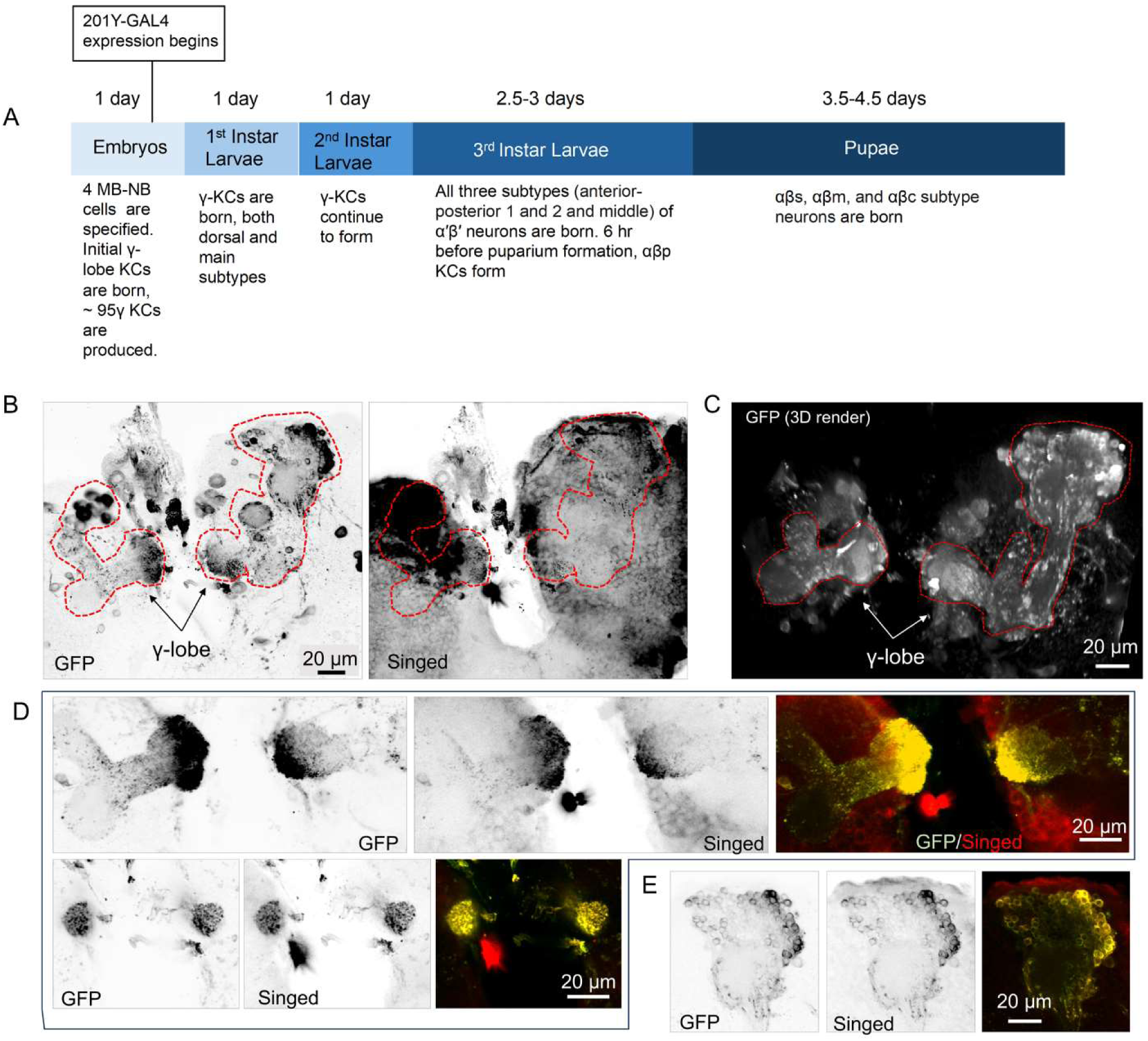
SN is widely expressed in the *Drosophila* larval brain and colocalizes with MB γ-lobe. **(A)** Timeline of MB development, highlighting the sequential birth of Kenyon cell (KC) subtypes. The 201Y-GAL4 driver begins expression during the embryonic stage and labels newly born γ-lobe KCs. **(B)** Representative z-stack maximum projection images of 3rd instar larval brains expressing UAS-mCD8:GFP under control of 201Y-GAL4, showing nascent γ-lobe axon tracts (left) and co-staining with anti-Singed (SN) antibody (right). Dashed red lines indicate the γ-lobes. **(C)** 3D-rendered image of the same genotype confirms early γ-lobe morphology marked by 201Y-GAL4. **(D)** Optical z-plane sections reveal co-localization of SN and GFP in γ-lobe neurons, consistent with SN expression in axons undergoing pathfinding. **(E)** SN is also detected in KC cell bodies, co-localizing with 201Y-driven GFP signal. Scale bars: 20 µm. Abbreviations: MB-NB, mushroom body neuroblast; KC, Kenyon cell; αβp, alpha and beta posterior; αβs, alpha and beta surface; αβm alpha and beta middle; αβc alpha and beta core.

**Figure S3:**
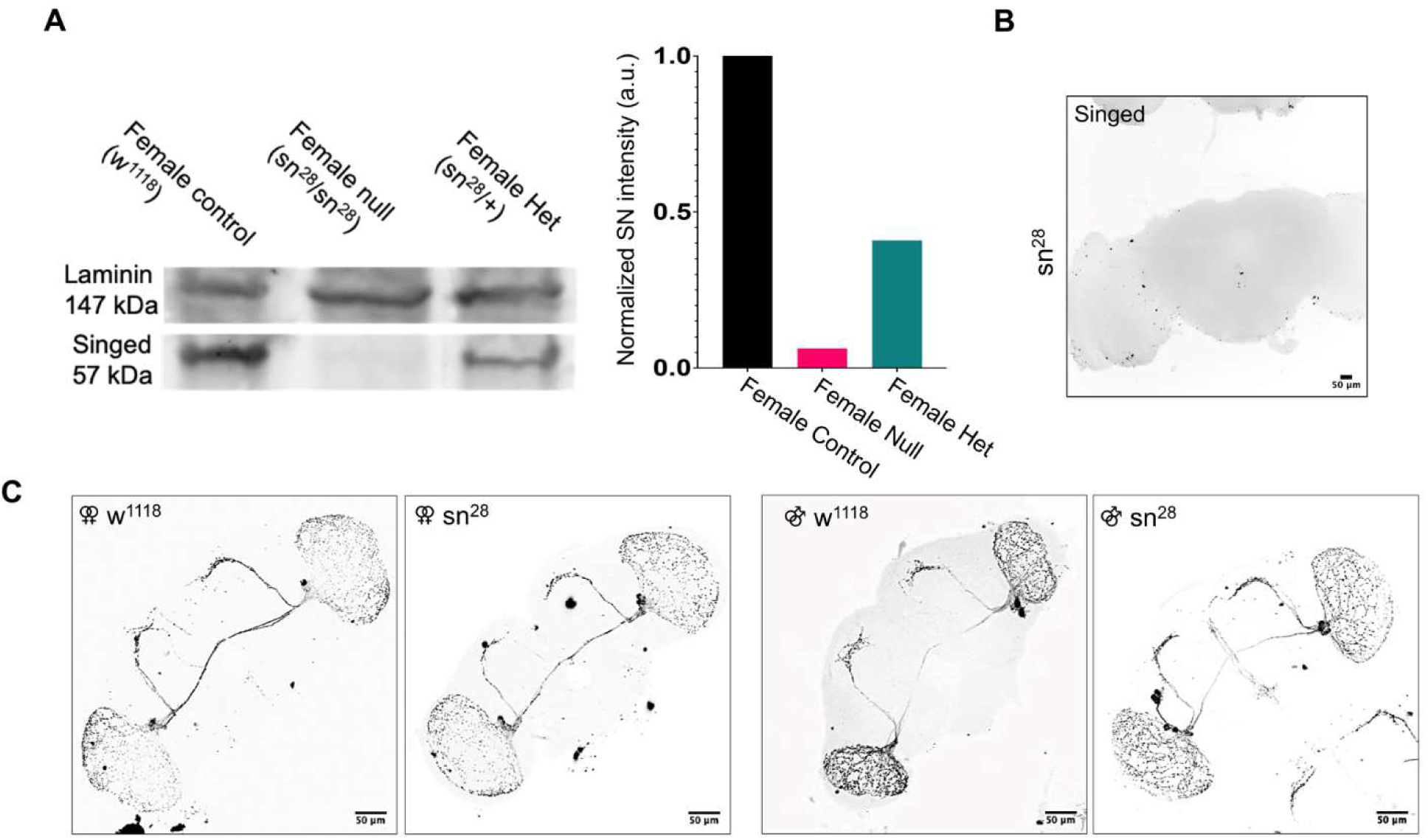
sn^28^ brains do not show any differences in PDF neuron projection compared to control w^1118^ brains. (**A**) Confirmation of singed knockout in sn^28^ line via western blot (n=1). Quantification of western blot via densitometry shows a reduction of SN in the heterozygous line and only background levels in the null line. (**B**) Brains dissected from sn^28^ flies and stained with anti-Singed antibody do not show any signal, further confirming that these flies are not producing SN protein. (**C**) Brains from female and male sn^28^ and w^1118^ flies were dissected and stained with an anti-PDF antibody. **We** do not observe any differences in PDF axon projection.

**Figure S4:**
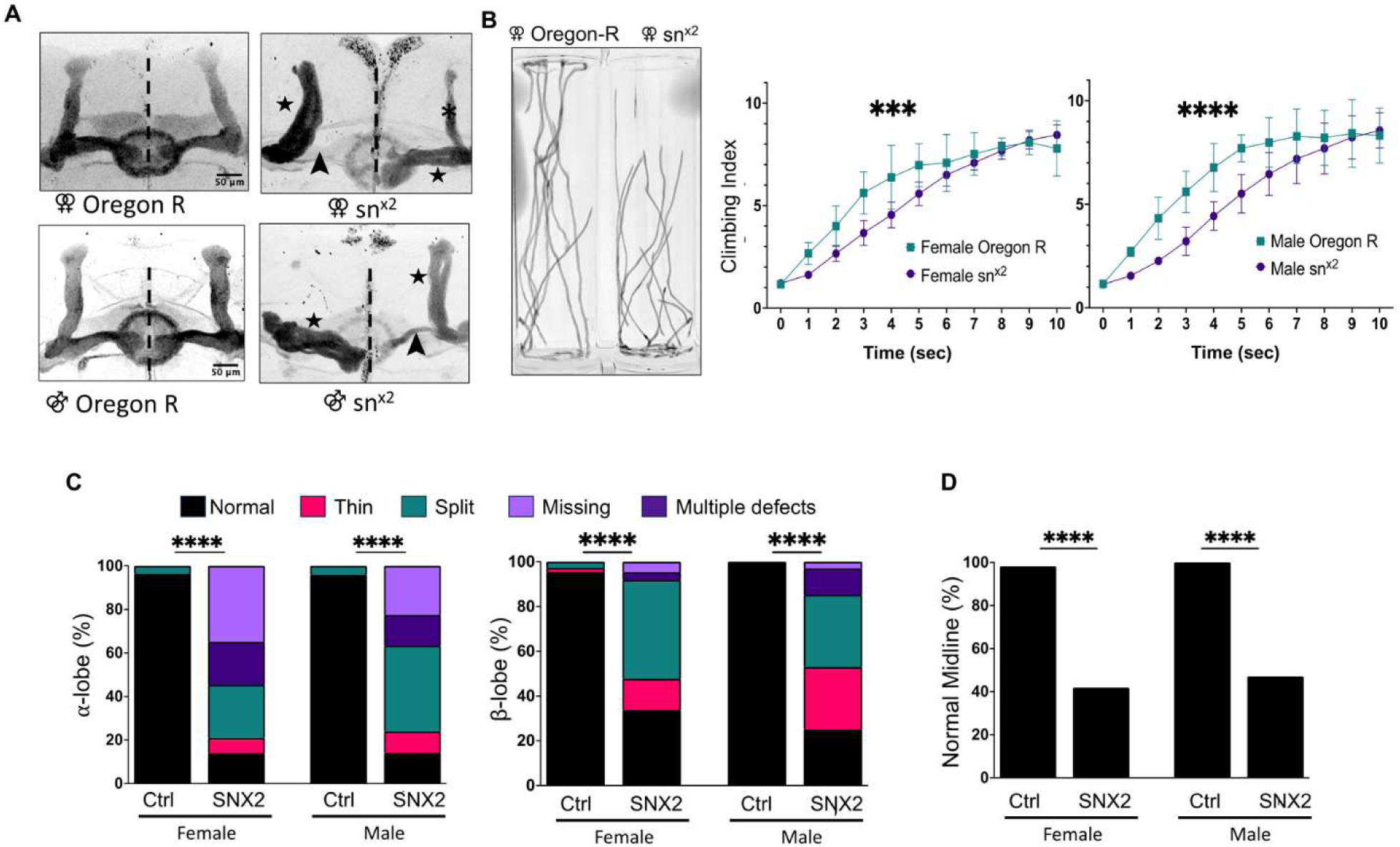
MB defects and NGA climbing deficits are also observed in another SN null line. (**A**) Representative images of Fas II staining in control female Oregon R and female sn^x2^ brains showing split ⍺-lobe, thin ⍺-lobe, split β-lobe, and a thin β-lobe (black arrowhead). Male Oregon R and male sn^x2^ brains also show similar defects to the female brains. female Oregon-R (n=54), female sn^x2^ (n=43), male Oregon-R (n=49), and male sn^x2^ (n=34). (**B**) Representative tracings of female Oregon-R and sn^x2^ flies 5 seconds after beginning to climb. Comparison of the climbing indices of female Oregon-R(n=54) and female sn^x2^ (n=40) flies shows a climbing deficit for the sn^x2^ flies, with the SN null flies taking longer to reach the top of the vial (analyzed using two-way ANOVA, p<0.0001). The same delay in reaching the top bin position is observed in male sn^x2^ (n=33) flies when compared to male Oregon-R (n=55), p<0.0001. (**C**) Quantification of ⍺- and β-lobe defects. Chi square analysis was used to calculate p-values, shown in these graphs is the analysis of total defects (**** indicates a p-value of <0.0001). (**D**) Quantification of midline defects observed in sn^x2^ brains (analyzed using Chi-square analysis, p<0.0001).

**Figure S5.**
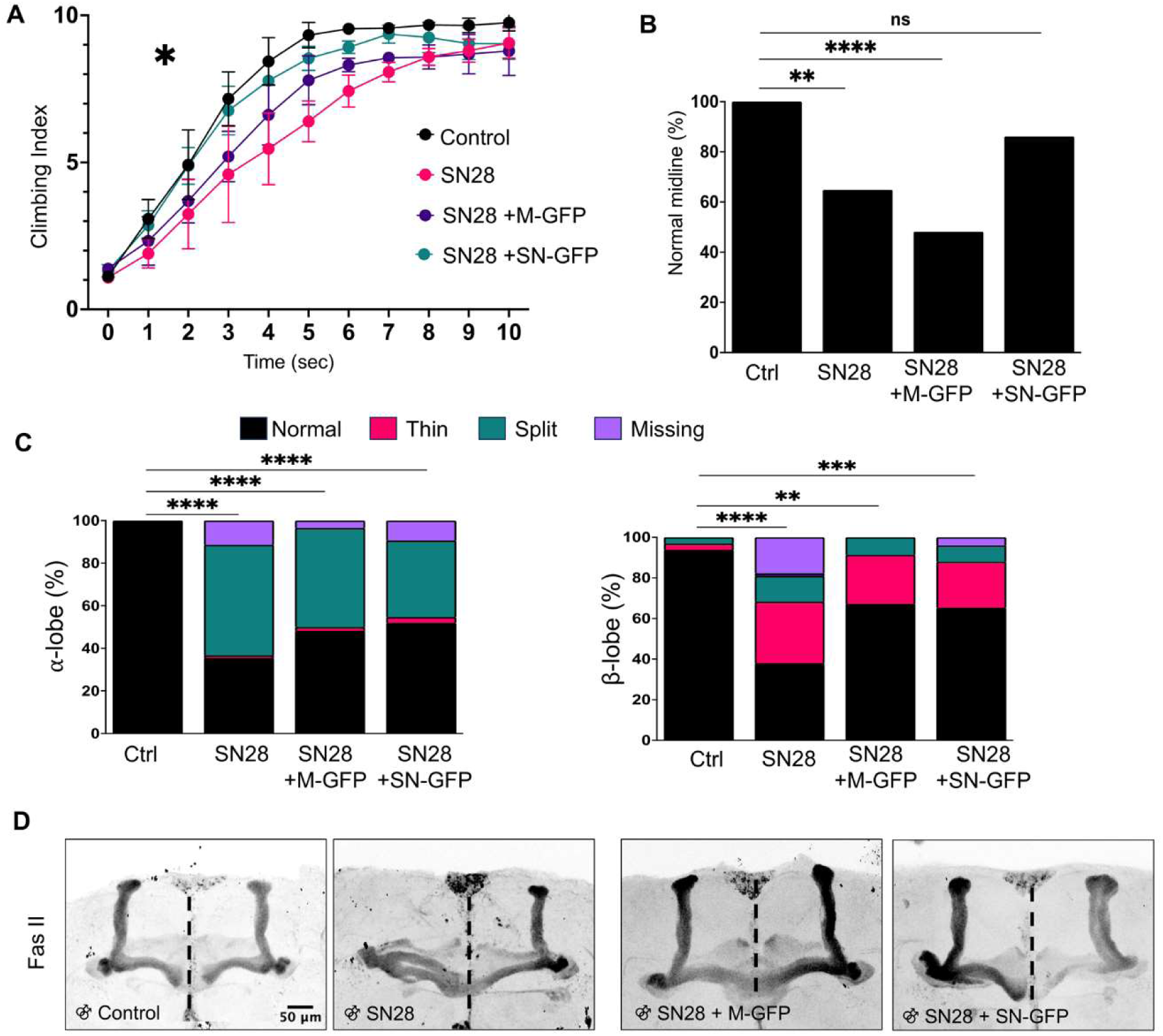
MB-specific expression of fascin1 is sufficient to rescue climbing and midline defects but not MB β-lobe and α-lobe defects in males. (A) Flies from 4 genotypes were collected and assessed using the NGA protocol. Flies expressing UAS-SN-GFP driven by 238Y-GAL4 are able to restore climbing ability to control w^1118^ levels (analyzed using two-way ANOVA, p=0.02. (B) MB-specific expression of UAS SN-GFP is sufficient to restore midline axon projection morphology in male flies. Statistical analysis was performed using Chi-square proportional analysis with post-hoc Marascuilo’s Procedure. Male w^1118^ n=32, male sn^28^ n=37 (p-value =0.0012), male sn^28^; 238Y-GAL4/UAS Myrs-GFP n=27 (p-value <0.0001), male sn^28^; 238Y-GAL4/UAS SN-GFP n=36 (p-value = 0.3255). (C) Expression of SN-GFP in the MB is unable restore MB lobe morphology. Brains were dissected and stained for Fas II. Quantification of MB ⍺- and β-lobe defects shows that there is not a rescue of MB lobe defects observed in sn^28^ flies upon MB-specific expression of WT GFP-Fascin1. Statistical analysis was performed using Chi-square proportional analysis with post-hoc Marascuilo’s Procedure. Male w^1118^ n=32, male sn^28^ n=37 (β-lobe p-value <0.0001, ⍺-lobe p-value <0.0001), male sn^28^; 238Y-GAL4/UAS Myrs-GFP n=27 (β-lobe p-value = 0.0099, ⍺-lobe p-value <0.0001), male sn^28^; 238Y-GAL4/UAS SN-GFP n=36 (β-lobe p-value = 0.0009, ⍺-lobe p-value <0.0001). (D) Representative images of MB ⍺- and β-lobes in the four genotypes examined.

### 3. Supplemental video 1-5

**Supplemental Video 1: GFP-fascin1 in the growth cone of primary hippocampal neurons.** Exogenously expressed GFP-fascin1 localizes to growth cone filopodia and can be seen dynamically coming on and off of the filaments. GFP-fascin1 can also be observed localizing to the lamellipodia in the growth cone as well.

**Supplemental Video 2: Wildtype mushroom body axons bifurcate and project vertically and medially.** Brains from control flies (Split-GAL4/UAS-myristoylated GFP; Split-GAL4/+) were dissected, PFA-fixed, and stained with anti-GFP and anti-Singed antibodies. 2-photon imaging generates a 3-D image of the axon projection from Kenyon cells and shows axons undergoing bifurcation into α and β lobe tracts which then follow guidance cues to form the α and β lobes.

**Supplemental Video 3: sn**^28^ **mushroom body axons bifurcate but appear to have guidance defects.** Brains from SN null flies expressing Split-GAL4/UAS-myristoylated GFP; Split-GAL4/+ were dissected, PFA-fixed, and stained with anti-GFP and anti-Singed antibodies. 2-photon imaging reveals that the axon tract extending from the Kenyon cells bifurcates but the α and β lobe tracts then do not follow proper guidance pathways. We observe midline crossing of the β lobes and additionally see the appearance of a split β lobe.

**Supplemental Video 4: sn**^28^ **mushroom body axons bifurcate but appear to have midline guidance defects.** As previously described, brains from SN null flies expressing Split-GAL4/UAS-myristoylated GFP; Split-GAL4/+ were dissected, PFA-fixed, and stained with anti-GFP and anti-Singed antibodies. When imaged, we see in this case a β lobe that appears to initially project medially but does not reach the midline and instead turns around. We also observe a β lobe that does project to the midline but some axons cross over the midline while the rest begin to turn vertically.

**Supplemental Video 5: sn**^28^ **flies show a locomotor defect.** Female w^1118^ flies are shown on the left and sn^28^ flies are on the right. When the flies are tamped to the bottom of the vial, the SN null flies show a delay in reaching the top when compared to control flies.

## Acknowledgements

We thank Drs. Kenneth Moberg and Dorothy Lerit for their help and guidance with the study. We sincerely thank Hengrui Zhang for his help with the primary hippocampal neuronal culture. This research is supported in part by a NRSA F31 fellowship from NIH-NINDS to KRH (NS127574), a R21 research grant from NIH-NINDS to JQZ (NS132393), and a Whitehall Foundation research grant to KRM. This work was also supported by the Emory University Emory Integrated Cellular Imaging Core Facility (RRID:SCR_023534).

## Author contributions

KRH led the effort in conducting most of the experiments and data analysis. ABP performed *Drosophila* brain dissections, immunostaining, imaging, and data analysis. SJ performed RNAi western blots and CAD cell imaging. CY helped with brain dissection and imaging. KRM helped with the experiments concerning cultured mammalian neurons and fascin manipulation. JQZ helped conceptualize the experiments, assisted with two-photon imaging, and provided guidance and constructive input to the study. KRH wrote the manuscript draft and worked with JQZ and KRM to finalize the manuscript.

